# Sialic acid ligands of CD28 block co-stimulation of T cells

**DOI:** 10.1101/2021.02.22.432333

**Authors:** Landon J. Edgar, Andrew J. Thompson, Vincent F. Vartabedian, Chika Kikuchi, Jordan L. Woehl, John R. Teijaro, James C. Paulson

**Author notes:** Correspondence to (J.C.P.).

## Abstract

Effector T cells comprise the cellular arm of the adaptive immune system and are essential for mounting immune responses against pathogens and cancer. To reach effector status, co-stimulation through CD28 is required. Here, we report that sialic acid-containing glycans on the surface of both T cells and APCs are alternative ligands of CD28 that compete with binding to its well-documented activatory ligand CD80 on the APC, resulting in attenuated co-stimulation. Removal of sialic acids enhances T cell activation and enhances the activity of effector T cells made hypofunctional via chronic viral infection through a mechanism that is synergistic with antibody blockade of the inhibitory PD-1 axis. These results reveal a previously unrecognized role for sialic acids in attenuation of CD28 mediated co-stimulation of T cells.

**One Sentence Summary:** Sialic acids attenuate the second signal required for T cell activation.

## Main Text

T cell-mediated immunity is central to host defense against pathogens, progression of autoimmunity, and elimination of cancer cells (*1*). To fully activate, T cells must receive two signals from an antigen presenting cell (APC) via a cell-cell interface called the immunological synapse (IS) (*2*). The first signal is antigen-specific, and is initiated by the T cell receptor (TCR) recognizing an antigenic peptide displayed on a major histocompatibility complex (MHC) on an APC (*3*). The second, ‘co-stimulatory’ signal is antigen-independent and mediated by CD28 on T cells, a receptor recruited to the IS by its protein ligands CD80/CD86, expressed on the APC (*4, 5*). Both signals are required for naïve T cells to differentiate into functional effector cells. In contrast to these activatory signals, T cell activation and continued function can be suppressed by immunological ‘checkpoint’ receptors such as PD-1 and CTLA-4. These inhibitory receptors on T cells are similarly recruited to the IS if their cognate ligands are expressed on the APC, leading to functional ‘exhaustion’ of T cells (*6*). Under these circumstances, while presented antigens are still recognized by the TCR, full activation cannot be achieved (*7*). Blocking checkpoint receptors using therapeutic antibodies has led to transformational developments in treatment of refractory cancers via functional revival of cancer-specific effector T cells from hypofunctional phenotypes (*8*). The success of these approaches has created enormous interest in understanding the detailed mechanisms that regulate activation of naïve and effector T cells.

Motivated by recent reports of the sialic acid-binding immunoglobulin-like receptors (siglecs) as immunological checkpoints (*9–11*), we took interest in work from several groups, starting nearly 40 years ago, that showed T cell activation was enhanced by enzymatic removal of sialic acids from the surface of T cells and/or APCs (*12–17*). Using an antigen-independent system, these seminal studies showed that T cell activation could be dramatically enhanced by prior treatment of syngeneic B cells with sialidases (*14*). This observation is mechanistically suggestive of a role of sialic acids in co-stimulation; however, no mechanism to explain this effect has been described. Here we show that CD28 recognizes sialylated glycans as alternative ligands that compete for binding to CD80. Further, we find that destroying sialic acid ligands leads to dramatic revival of hypofunctional PD-1^+^ T cells, and that this enhancement is synergistic with blockade of the PD-1 checkpoint inhibitory pathway.

To analyze the impact of sialic acids on antigen-specific T cell activation we used chicken ovalbumin (OVA)-specific CD4^+^ and CD8^+^ T cells (OT-I and OT-II cells respectively) in combination with bone marrow-derived dendritic cells (DCs) matured with bacterial lipopolysaccharide (LPS). Treatment of T cells and DCs with sialidase from *Vibrio cholerae* efficiently removed cell surface sialic acids from the most common NeuAcα2-3Gal and NeuAcα2-6Gal linkages found on glycoproteins of T cells and DCs, as detected by fluorescent lectin staining (Fig. 1A). In the context of T cell activation, sialidase-treated co-cultures of OVA-presenting DCs with OT-I or OT-II cells lead to a significant enhancement in clonal expansion of both T cells types, as measured by dilution of the proliferation reporter dye CellTrace Violet (CTV) (Fig. 1, B–D). This effect was conserved under treatment with an alternative sialidase from *Streptococcus pneumoniae*, and when alternatively matured DCs or splenocytes were substituted as APCs (fig. S1 and fig. S2). We observed no effect when heat denatured enzyme (Δ) was used, confirming the dependence on specific sialidase activity.

**Fig. 1.**
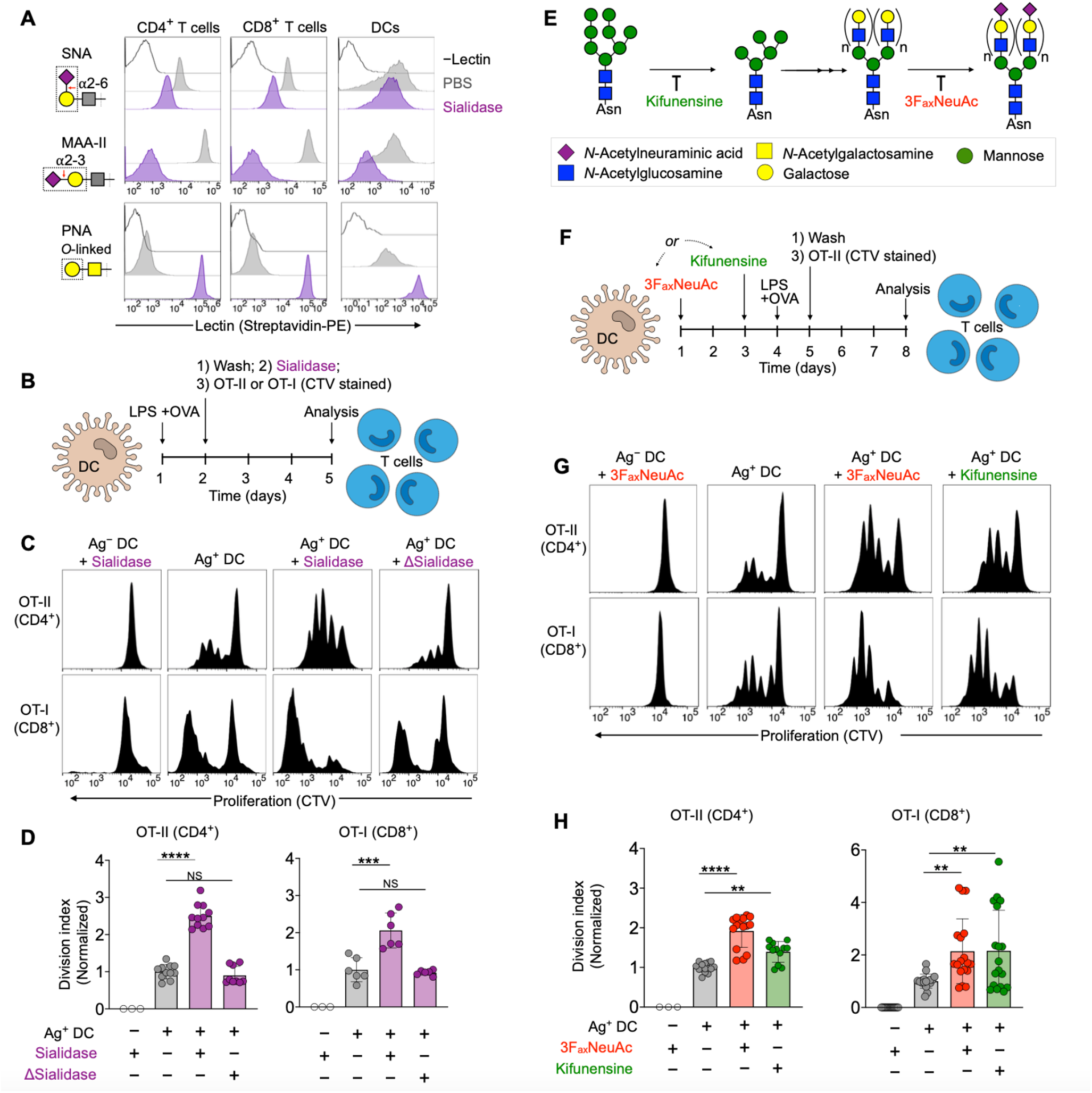
Desialylation enhances antigen-dependent activation of T cells. (**A**), Lectin staining of T cells (from a mixture of splenocytes) and bone marrow-derived dendritic cells before (PBS) and after treatment with sialidase from *V. cholerae* (55 mU). Red arrow indicates site of action of sialidase and dashed box shows lectin recognition motif. (**B**), Antigen-specific T cell expansion assay setup. (**C**), T cell proliferation histograms (dilution of CTV) for co-cultures in the presence of sialidase from *V. cholerae*. DC to T cell ratio was 1:2. Δ = heat inactivated. (**D**), Quantification of T cell expansion data from c (*n* ≥ 3). (**E**), Schematic of glycosylation pathways inhibited by 3F_ax_NeuAc and kifunensine. See Fig. 2d for pictogram definitions of monosaccharides. (**F**), Expansion of T cells using DCs selectively desialylated via pretreatment with 3F_ax_NeuAc or kifunensine. (**G**), T cell proliferation histograms for co-cultures as set up in f. DC to T cell ratio was 1:2. (**H**), Quantification of T cell expansion data from g (*n* ≥ 3). Mean ± s.d. (**D, H**). \*\**P* ≤ 0.01, \*\*\**P* ≤ 0.001, \*\*\*\**P* ≤ 0.0001, *ns* = not significant (**D, H**). One-way ANOVA followed by Tukey’s multiple comparisons test (**D, H**). Normalized division index corresponds to T cell division index for sialidase-treated cultures divided by the division index for the corresponding PBS treated control. T cell proliferation data is pooled from at least 3 separate experiments.

To assess the impact of selective desialylation on APCs, we cultured DCs in the presence of two specific inhibitors of glycosylation, kifunensine or 2,4,7,8,9-pentaacetyl-3F_ax_-Neu5Ac-CO_2_Me (3F_ax_NeuAc) (*18*). Kifunensine acts early in N-linked glycan maturation, preventing further processing of precursor high-mannose type glycans (and thus subsequent addition of sialic acids as terminal sugar residues) (*19*). In contrast, 3F_ax_NeuAc directly prevents sialic acid transfer through inhibition of sialyltransferases (Fig. 1E) (*18–20*). Both kifunensine and 3F_ax_NeuAc were able to reduce DC sialylation on the timescale of T cell activation assays, albeit to a lesser extent than constant exposure to sialidase (fig. S3). Nonetheless, DCs treated with either inhibitor were significantly more potent in activating normally sialylated CD4^+^ and CD8^+^ T cells as compared to untreated controls, suggesting that sialic acids on N-glycans within the glycocalyx of DCs suppress activation of T cells (Fig. 1, F–H). To determine if this effect was dependent on IS formation with the APC or was mediated by secreted soluble factors, we conducted experiments where T cells and DCs were physically separated using a transwell system. Neither kifunensine nor 3F_ax_NeuAc-desialylated DCs showed any ability to enhance T cell activation over PBS controls when physically separated from untreated T cells, and expression of both activatory / inhibitory receptors on the DCs remained unchanged (fig. S4 and fig. S5). Taken together, these data suggested that sialic acids on *N*-linked glycans negatively impact signaling between T cells and APCs, and that sialic acids of APCs can contribute to attenuation of antigen-mediated T cell activation through direct interference at the IS.

The co-stimulatory receptor CD28, its ligands, CD80/CD86, and the inhibitory receptors PD-1/PD-L1/PD-L2/CTLA-4 are all immunoglobulin (Ig)-like cell surface receptors. We therefore considered the possibility that one or more of these proteins might directly bind sialic acids as ligands, similar to the sialic acid-binding immunoglobulin-like lectin (Siglec) family of receptors, which are also members of the Ig-like superfamily (*21–26*). Indeed, like the Siglecs, these proteins all possess an N-terminal V-set Ig domain and share significant sequence and structural homology with the sialic acid ligand-binding region present in each siglec (Fig. 2A, top, and fig. S6 and fig. S7) (*21–26*). To evaluate potential sialoside-binding activity, we employed recombinant chimeras comprising N-terminal V-set Ig domains (and C2-set domains where appropriate) fused to the Fc-domain of human IgG. These were applied to glycan arrays that contained a diverse library of glycans capped with sialic acids including intact *N*-linked and *O*-linked glycans, as well as fragments representing terminal sequences commonly found in glycoprotein and glycolipid glycans. Representative glycans without terminal sialic acids were included as controls (Fig. 2B, table S1) (*27*). Importantly, human and murine CD28-Fc exhibited strikingly similar binding profiles to sialoglycans on the array, including several *N*-linked and *O*-linked glycans with the sequence NeuAcα2-3/6Galβ1-4GlcNAc, and to shorter fragments with an additional sulfate on the Gal or GlcNAc (Fig. 2, C and D). Quantitative binding to sialosides was also evaluated via surface plasmon resonance (SPR), revealing that recombinant monomeric CD28 binds to a representative glycan (#19, see table S1) with *K*_d_ = 112 μM (Fig. 2E). In contrast, none of the other Fc-chimeras examined for PD-1, PD-L1, PD-L2, CDLA-4, CD80 or CD86 displayed significant binding to the array (Fig. 2C and fig. S8). Importantly, when murine and human CD28-Fc were pre-complexed with their respective CD80 ligands prior to exposure to the array, binding to sialic acids was blocked (Fig. 2C). These data reveal that sialylated glycans are alternative ligands for both human and murine CD28, and that binding to CD80 appears competitive with binding to sialosides (Fig. 2A, bottom).

**Fig. 2.**
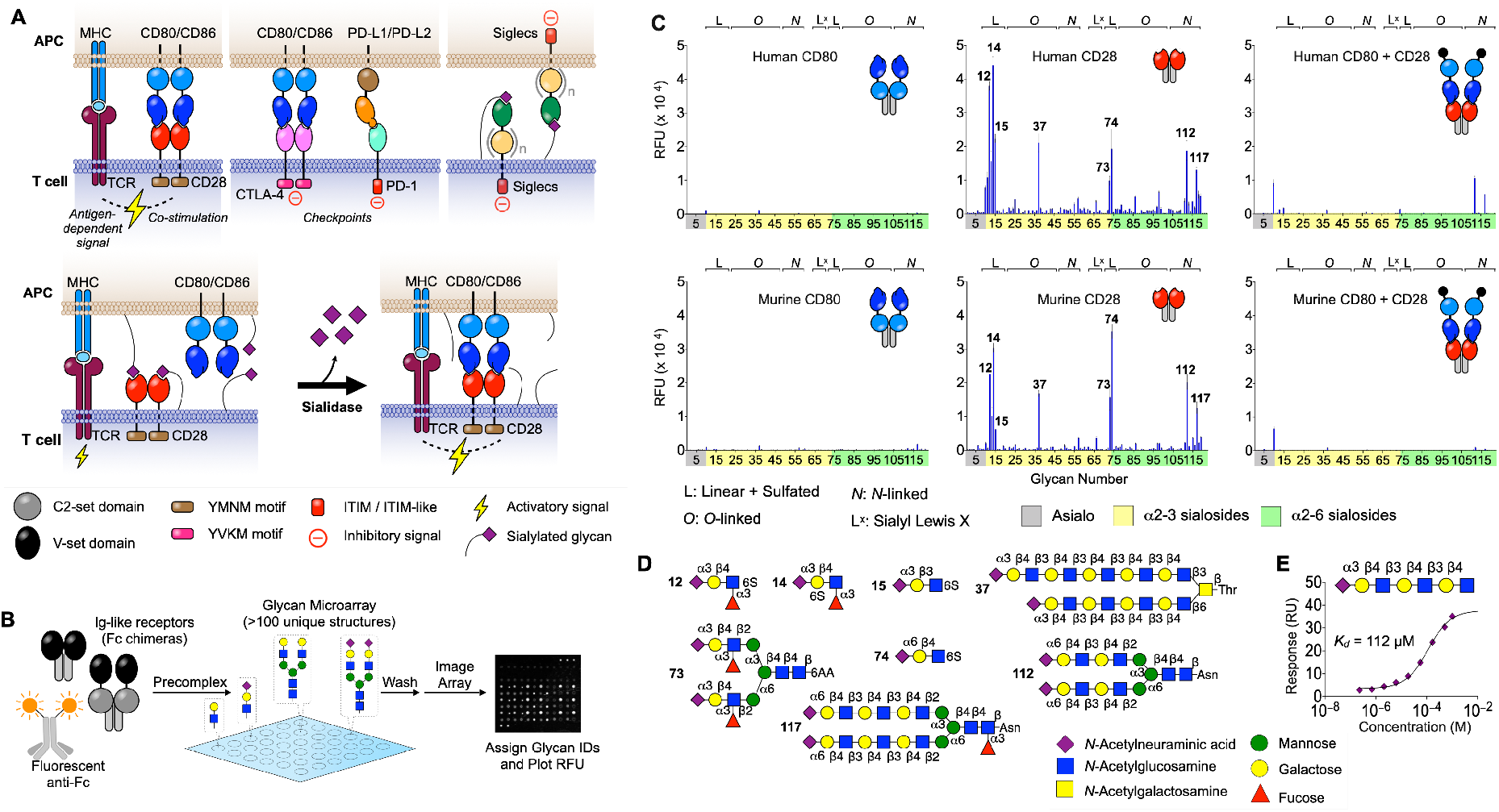
CD28 binds sialylated glycans. (**A**), Survey of Ig-like receptors at the IS (top). Competition between sialic acid ligands of CD28 and CD80 at a normally-sialylated IS and increased costimulatory interactions as a result of treatment with sialidase (bottom). (**B**), Glycan array screening workflow for IS receptors. (**C**), Glycan array binding data for human and mouse CD80, CD28, and both receptors pre-complexed. Compound IDs for top hits are indicated on the plots for CD28 (*n* = 4 for each peak). (**D**), Structures of glycan ligands for CD28. (**E**), Steady state binding of α2-3 sialyl-diLacNAc to surface immobilized CD28-GFP measured via SPR. Black line represents steady-state fit. Mean ± s.d (**C**).

To further assess the impact of sialic acids on CD28:CD80 interactions we measured direct binding of CD28-Fc to DCs and CD80-Fc to T cells, with and without prior sialidase treatment (Fig. 3). Binding of CD28-Fc to desialylated DCs was significantly enhanced compared to untreated cells and could be blocked in the presence of anti-CD80, demonstrating that sialic acids can act in *‘trans’* on the DC to reduce binding of CD28 to CD80 (Fig. 3, A and B). Similarly, binding of CD80-Fc to either CD4^+^ or CD8^+^ T cell populations was dramatically enhanced by sialidase treatment (Fig. 3, C and D). These results suggest that sialic acid-containing glycans present on the surface of T cells may also act in *‘cis’* as competitive ligands, sequestering available CD28 and thus inhibiting binding to CD80 (Fig. 3C). In this context, it is notable that the sialic acid content in the lymphocyte glycocalyx is >100 mM, (*28*) far above the likely *K*_d_ for binding of most sialic acid ligands to CD28 (Fig. 2E). Although sialic acid ligands on the T cell block CD80-Fc binding, the sialoside:CD28-Fc interaction is not sufficiently avid to support binding of CD28-Fc to T cells alone, with staining appearing identical to sialidase-treated controls (fig. S9). Taken together, these results frame a role for sialic acids in attenuating T cell co-stimulation since binding of CD28 to CD80 is required for productive co-stimulation (Fig. 2A) (*4, 5*).

**Fig. 3.**
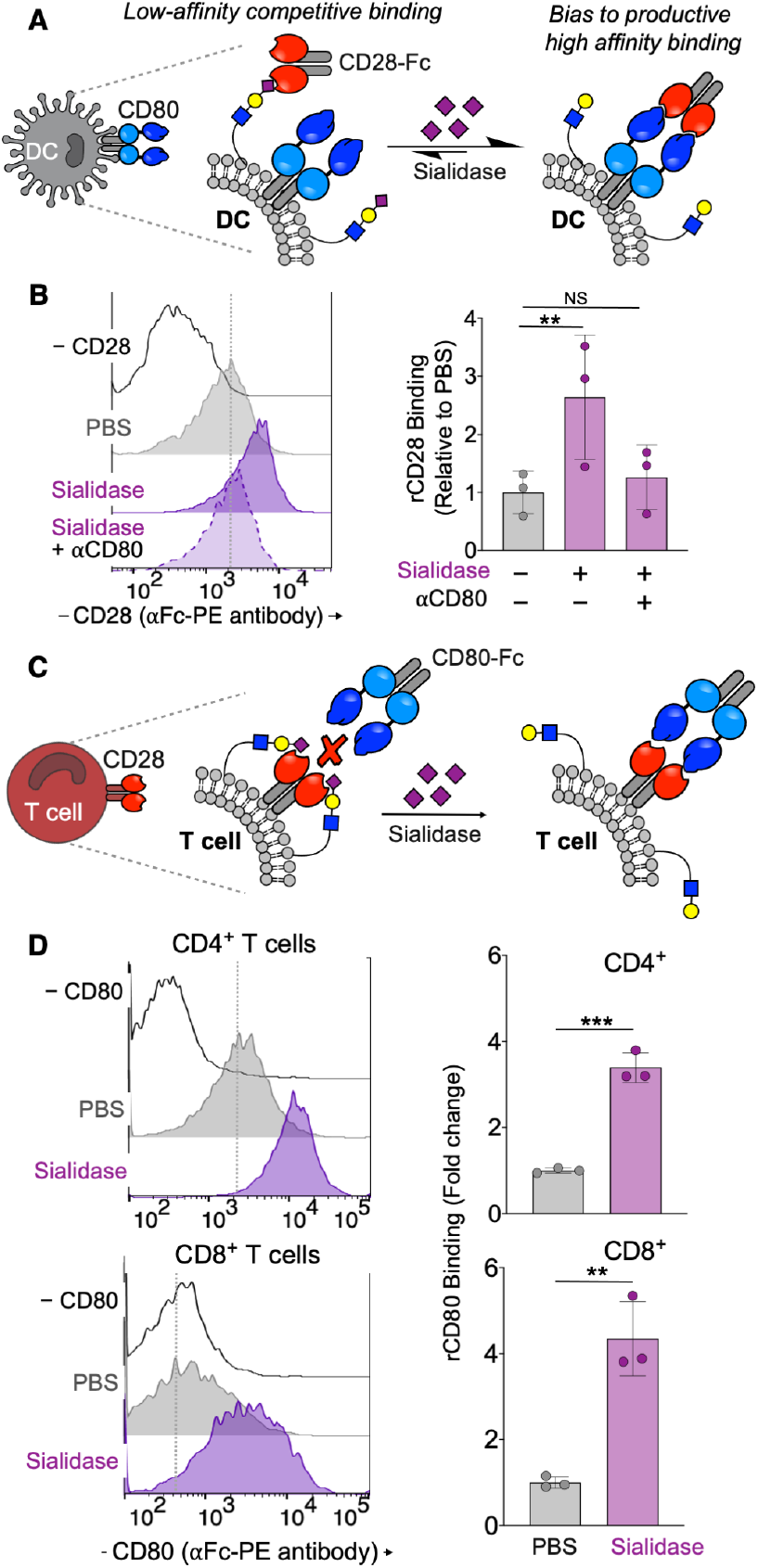
Sialylated glycans on T cells and DCs impair CD28 binding to CD80. (**A**), Schematic of the impact of DC desialylation to costimulatory synapse formation. (**B**), Staining of bone marrow-derived DCs with recombinant CD28-Fc. Sialidase from *V. cholerae* (55 mU) was used to desialylate DCs. (**C**), Schematic of the impact of T cell desialylation to costimulatory synapse formation. (**D**), Staining of splenic T cells from WT mice with recombinant CD80-Fc. Sialidase from *V. cholerae* (55 mU) was used to desialylate T cells. Mean ± s.d (**B, D**). \*\**P* ≤ 0.01, \*\*\**P* ≤ 0.001, ns = not significant (**B, D**). One-way ANOVA followed by Tukey’s multiple comparisons test as a paired analysis (**B, D**).

Since co-stimulation through CD28 is key to reviving hypofunctional T cells through checkpoint blockade therapy (*29*), we reasoned that hypofunctional T cells could be revived more efficiently by removal of sialic acids. To test this, we used leukocytes from mice infected with lymphocytic choriomeningitis virus (LCMV, clone 13) as a source of hypofunctional (PD-1^+^) polyclonal CD8^+^ T cells with defined antigen specificity (Fig. 4, A and B) (*30*). Splenocytes from these animals were cultured ex vivo for 5 h in the presence of viral peptide antigen (immunodominant peptide from viral glycoprotein (residues 33–41, GP33) or subdominant peptide from viral nucleoprotein (residues 205–212, NP205)) and/or sialidase (Fig. 4A). Sialidase treatment increased the percentage of functional CD8^+^ T cells over GP33 alone, as defined by increased expression of both granzyme B (GrnzB) and interferon-*γ* (IFN-*γ*) (Fig. 4, C and E). Importantly, we observed that hypofunctional (PD-1^+^) CD8^+^ T cells also exhibited increased activation when treated with sialidase and re-stimulated with either GP33 or NP205 (Fig. 4, C and E). No significant activation was observed in the absence of antigen with or without sialidase. To place these findings into context with direct blockade of the PD-1 axis, we repeated the re-stimulation assay with anti-PD-L1 (αPD-L1) antibody blockade and/or sialidase at an extended time point (72 h) (Fig. 4F). We chose αPD-L1 to facilitate comparisons with previous studies using similar systems (*29, 31*). Here, splenocytes stimulated with GP33 and sialidase produced more IFN-*γ*’ as compared to antigen alone, with an effect comparable to αPD-L1 treatment (Fig. 4G). Importantly, combination treatment with both αPD-L1 and sialidase resulted in the strongest and most significant reactivation of CD8^+^ T cells for both GP33 and NP205 peptides (Fig. 4G). This result is consistent with our expectation that enhanced co-stimulation as enabled by enzymatic desialylation can enhance the efficacy of checkpoint blockade strategies.

**Fig. 4.**
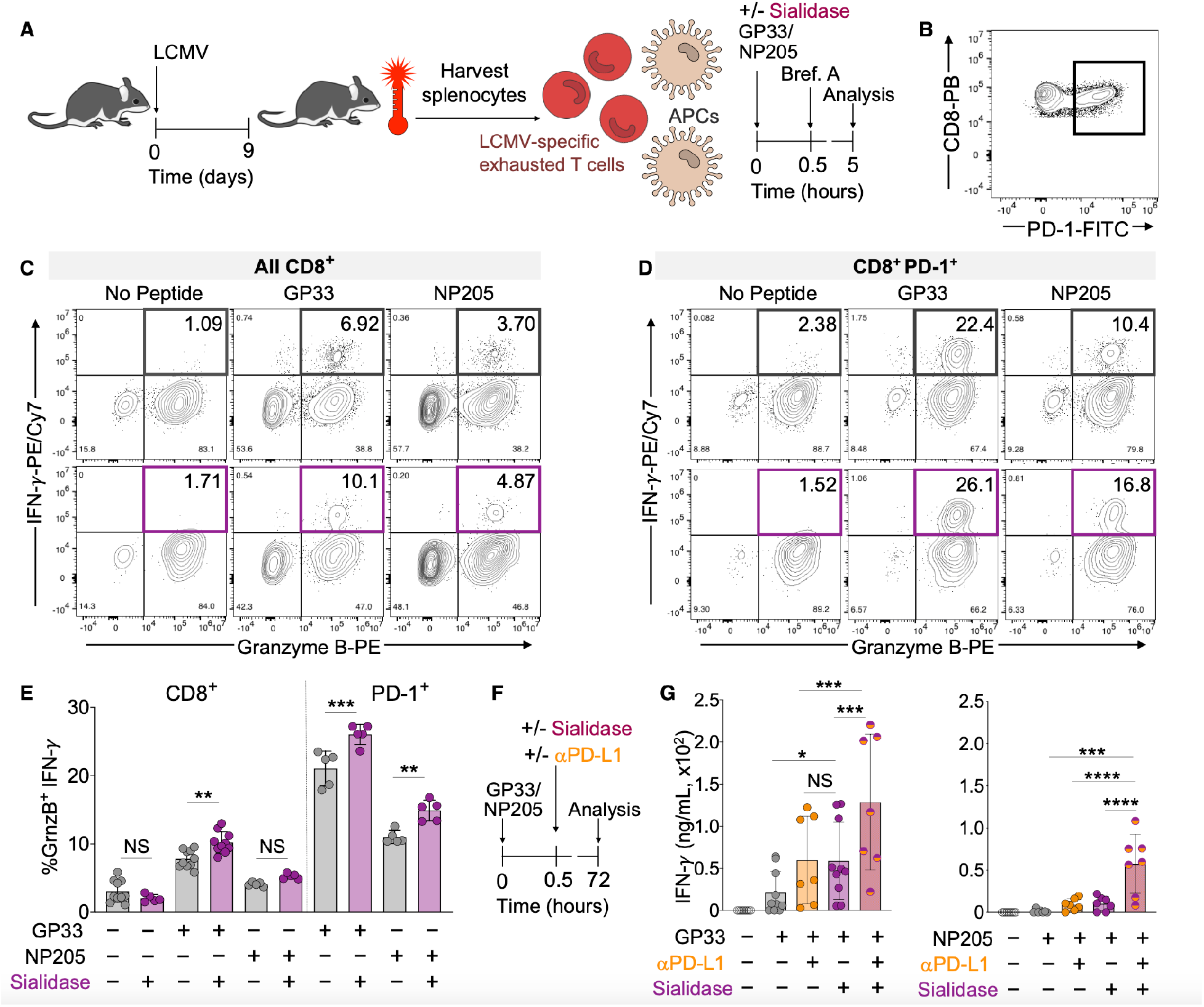
Sialidase enhances revival of functionally hypofunctional T cells. (**A**), Generation of polyclonal hypofunctional LCMV-specific CD8^+^ T cells from WT C57BL6/J mice. Animals were infected with LCMV (2×10^6^ pfu, clone 13). Spleens were harvested on day 9 post-infection and splenocytes (containing a mixture of leukocytes including T cells and APCs) were cultured in the presence of GP33 or NP205 LCMV peptide antigens and/or sialidase from *V. cholerae* (55 mU). Brefeldin A (Brf. A) was added at 0.5 h and cytokine production in CD8^+^ T cells was assessed at 5 h via flow cytometry. (**B**), Representitive PD-1 expression on CD8^+^ T cells from LCMV infected mice. (**C**), Representative density maps of activated (GranzymeB^+^ IFN-*γ*^+^) LCMV antigen-specific polyclonal CD8^+^ T cells. (**D**), Representative density maps of activated PD-1^+^ (GranzymeB^+^ IFN-*γ*^+^) LCMV antigen-specific polyclonal CD8^+^ T cells. (**E**), Quantification of the percentage of activated CD8^+^ T cells from c and d (*n* ≥ 5). (**F**), Assay workflow for longer term (72 h) activation of T cells made hypofunctional via chronic LCMV infection as in a. g, Quantification of antigen-induced IFN-*γ* production by polyclonal CD8^+^ T cells from LCMV infected mice as in a and f. IFN-*γ* was quantified via ELISA after 72 h *ex vivo* stimulation with antigen and anti-mouse PD-L1 (25 μg/mL) and/or sialidase from *V. cholerae* (55 mU) (*n* ≥ 4). Mean ± s.d (**e, g**). \**P* < 0.05, \*\**P* ≤ 0.01, \*\*\**P* ≤ 0.001, \*\*\*\**P* ≤ 0.0001, ns = not significant (**E, G**). One-way ANOVA followed by Tukey’s multiple comparisons test as a paired analysis (**E, G**).

Co-stimulation through CD28 is an indispensable signal required for full activation of naïve and effector T cells — a role that has been recognized for over 30 years (*32*). Here we have shown that CD28 recognizes sialic acid-containing glycans as ligands, and that CD28-glycan complexes have a reduced capacity to interface with canonical activatory ligands expressed by APCs. We propose that sialic acid-mediated attenuation of CD28:CD80 interactions provides a mechanistic basis for the decades-old observation that sialidase treatment of T cells or APCs enhances antigen-specific T cell activation. We and others have shown that sialic acid-containing glycans on T cells are dynamically remodeled during differentiation and activation as a result of altered expression of sialyltransferase and neuraminidase genes (*33, 34*). When placed into context with this work, these observations suggest that such remodeling is a biologically authentic step in T cell activation and could tune the amount of CD28 available for co-stimulation through altered expression of sialic acid ligands in *cis*. Similarly, various APCs (e.g. DCs, B cells, cancer cells, etc.) have cell-type-specific glycosylation with unique compositions of sialic acid-containing glycans that could act to attenuate co-stimulation in *trans*. Therapeutic precedence for disruption of sialic acids in the latter case was shown by Gray et al., where sialidase-antibody conjugates selectively desialylate tumor cells to destroy ligands for inhibitory siglecs expressed on natural killer cells (*10, 11*). This strategy profoundly suppressed tumor growth, and is also mechanistically consistent with enzymatic removal of sialic acids that inhibit CD28-mediated co-stimulation of T cells. In addition, others have demonstrated that reducing sialylation on murine melanoma tumor cells promotes generation of effector T cells and slows disease progression (*35*). Our discovery that sialidase can enhance reactivation of hypofunctional T cells, and that this effect is synergistic with blockade of the PD-1 axis, is also relevant to these and other disease models where PD-1 is expressed on tumor-infiltrating lymphocytes (*36*). Thus, approaches to reduce the sialic acid content of T cells or APCs may have value in settings requiring enhanced co-stimulation via CD28 for generation of effector or revival of hypofunctional T cells, especially in scenarios where blockade of the PD-1 axis alone is insufficient.

## Acknowledgments

The authors thank Cory Rillahan for preparing 3F_ax_NeuAc, Charli Worth for assistance with acquiring glycan microarray data, Britni Arlian Cruz and Jasmine Stamps for genotyping mice, Alan Saluk and Brian Seegers for assistance with flow cytometry, and Haissi Cui and Corwin Nycholat for helpful discussions.

## Author contributions

LJE and JCP designed the study. LJE, AJT, VFV, CK, and JLW executed experiments. JRT advised on LCMV-related experiments. The manuscript was written by LJE and JCP with input from all authors.

## List of Supplementary Materials

Materials and Methods

Figs. S1 to S9

Table S1 Reference (*37–40*)

## Data and Materials Availability

Glycan array data in MIRAGE format is available.

