## Supporting Information for "Sialic acid ligands of CD28 block co-stimulation of T cells"

**This PDF file includes:**

Materials and Methods
Figs. S1 to S9
Table S1

**Materials and Methods:**

**Bone marrow-derived dendritic cells.** To culture bone marrow derived dendritic cells (DCs), femurs from male WT C57BL/6J mice were harvested and crushed in HEPES buffer (20 mM) using a mortar and pestle. The resultant cell suspension was filtered through nylon mesh (40 µm) and diluted with complete media (RPMI, 10% fetal bovine serum (FBS), 1% penicillin/streptomycin) to achieve a final cell concentration of $2x10^6$ /mL. GM-CSF (BioLegend) or Flt3L (BioXcell) was then added to final concentrations of 20 and 200 ng/mL respectively. For GM-CSF cultures, additional GM-CSF (final concentration = 40 ng/mL) was added on day 3 of culture. After 7 days, cells were collected, centrifuged (350 rcf), and washed with complete media. To prepare DCs for stimulation of OT-I/OT-II cells, OVA (0.5 mg/mL) (Worthington Biochemical Corp.) and LPS (100 ng/mL) (Sigma-Aldrich) was added on day 6 of culture. For treatment with inhibitors of glycan biosynthesis, Kifunensine (25 µM) or $3F_{ax}$NeuAc (200 µM) were added on day 6 or day 4 of culture respectively.

**T cell expansion assays.** Splenocytes from $Ly5a^{+/+}$ OT-I (C57BL/g-Tg(TcraTcrb)1100Mjb/J) or $Ly5a^{+/-}$ OT-II (B6.Cg-Tg(TcraTcrb)425Cbn/J) mice were stained with the proliferative reporter dye cell trace violet (CTV) as previously reported (*37*). $CD8^+$ or $CD4^+$ TCR $V\alpha2^+$ cells were quantified via flow cytometry using a portion of the CTV stained splenocytes. Antigen loaded DCs were then added to cultures with CTV stained splenocytes at a ratio of 1 DC : 2 OT-I or OT-II cells (as quantified in previous step, $2x10^4$ DC and $4x10^4$ OT-I/II cells) in tissue cultured-treated polystyrene U-bottomed 96 well plates. For sialidase treatments, enzyme was added during this setup stage. After 3 days in culture (37 °C, 5% $CO_2$), cells were prepared for analysis via flow cytometry.

**Binding of Fc-chimeras and plant lectins to DCs and T cells.** DCs were prepared as described above (matured with LPS). Splenocytes from WT C57BL/6J mice were used as a source of both $CD4^+$ and $CD8^+$ T cells. DCs or splenocytes were treated with sialidase from *V. cholerae* (55 mU) for 45 min. at 37 °C followed by washing with PBS. DCs were then stained with unlabeled anti-mouse CD16/32 (Fc block) followed by fluorescent antibodies against CD11c (clone N418, BioLegend) and MHC II (I-A/I-E, clone M5/114.15.2, BioLegend). Similarly, splenocytes were first Fc blocked and then stained with fluorescent antibodies against CD3 (clone OKT3, ThermoFisher), CD4 (clone GK1.5, BioLegend), CD8a (clone 53-6.7, BioLegend), and CD19 (as a lineage gate to exclude B cells, clone 6D5, BioLegend). Fc blocked and antibody-stained cells were then stained with either plant lectins or recombinant CD28-Fc/CD80-Fc. For staining with plant lectins, biotinylated SNA (Vector Laboratories), MAA-II (Vector Laboratories), PNA (Vector Laboratories), or ECA (Vector Laboratories) were first precomplexed with fluorescent streptavidin for ~30 min. at 4 °C and subsequently incubated with DCs or splenocytes for 20 min. at 4 °C. Cells were then washed twice and analyzed via flow cytometry. For staining with recombinant CD28-Fc or CD80-Fc (BioLegend), DCs or splenocytes were incubated with recombinant proteins (4 μg/mL) for 20 min. at 4 °C, washed twice, and subsequently stained with fluorescent goat anti-human Fc (polyclonal, Jackson ImmunoResearch Laboratories). Cells were then washed twice and analyzed via flow cytometry.

**Recombinant *Vibrio cholerae* sialidase (nanH).** A plasmid harboring a fragment of the native *V. cholerae* genome including the nanH gene (pCVD364 (*38*)) was obtained as a gift from James Kaper and distributed via Addgene (plasmid # 63739 ; http://n2t.net/addgene:63739 ; RRID:Addgene_63739). The full nanH gene, minus a 50-amino acid fragment at the N-terminus, was amplified by PCR and subcloned via HiFi assembly (NEB) into a modified pET23 backbone using the following primers: FWD: 5'-TTT AAG AGG AGG TCA TAT GGC ACT TTT TGA CTA TAA CG-3', REV: 5'-AAT GGT GAT GGT GAT GAT GGT TTT GCG ATA AGG TCA-3'; resulting in direct fusion of a $His_6$ purification tag to the C-terminus.

For expression, pET23-nanH was transformed into *E. coli* BL21 (DE3) electrocompetent cells and grown in approximately 1.5 L of LB growth media to a final $OD_{600nm}$ of 0.8. Gene expression was induced through addition of 0.2 mM (final) exogenous IPTG and cultures were incubated overnight at 16 $^{o}$C prior to purification. Neuraminidase-expressing *E. coli* cells were harvested by centrifugation at 4500 xg, resuspended in cold 30 mM MES pH 6.5, 300 mM NaCl (approximately 15 mL per gram of cells), and lysed for purification via sonication. Clarified bacterial supernatant (post centrifugation at 15,000 xg) was passed through 0.45 µm syringe filters and loaded onto a 5mL NiNTA FF crude column (GE), preequilibrated in 30 mM MES pH 6.5, 300 mM NaCl, for $Ni^{2+}$-affinity purification.

For endotoxin removal, column-bound neuraminidase was washed with approximately 75 column volumes of cold 30 mM MES pH 6.5, 300 mM NaCl, 0.1% triton X-114 at 4 $^{o}$C. The column was then re-equilibrated in approximately 50 column volumes 30 mM MES pH 6.5, 300 mM NaCl prior to elution. Bound neuraminidase was eluted in a 30 mL gradient, exchanging into 30 mM MES pH 6.5, 300 mM NaCl, 500 mM imidazole. Protein-containing fractions were identified and purity determined via SDS-PAGE analysis. Pure neuraminidase-containing fractions were pooled, concentrated via centrifugation using a 50 kDa MWCO ultrafiltration device (Amicon), and applied to a Superdex 200 (16-600; GE) column preequilibrated in 30 mM MES pH 6.5, 300 mM

NaCl for further size-exclusion chromatography purification. Protein-containing fractions were again analyzed and purity determined via SDS-PAGE analysis. Final fractions corresponding to a single size-exclusion peak were concentrated to approximately 2.5 mg $mL^{-1}$ (final), aliquoted, snap frozen in liquid $N_2$, and stored at -80 $^{o}$C.

**Sialidase activity determination.** Activity of pure *V. cholerae* neuraminidase stock was determined via standard 4-methylumbelliferyl-α-D-N-acetylneuraminic acid (MUNANA) assay. Briefly, stock enzyme aliquots were thawed and diluted in 10-fold series ranging from $10^{-2}$ to $10^{-7}$ (25 µg $mL^{-1}$ to 250 pg $mL^{-1}$ final) and applied to individual 100 µL reactions in a pre-warmed 96-well plate containing 30 mM MES pH 6.0, 300 mM NaCl, 1 mM $CaCl_2$, 1% BSA, 100 µM (final) MUNANA. Following enzyme addition, reactions were incubated at 37$^{o}$C, with 10 µL time-point samples removed at the following intervals: 2, 5, 10, 20, 40 min, and rapidly mixed with 90 µL quenching solution (0.15 M $Na_2CO_3$, 0.15 M glycine pH 10) pre-aliquoted into a black 384-well plated. Upon completion, 4-methylumbelliferyl release for all samples was determined by fluorescent readout ($\lambda_{ex.}$ = 365 nm, $\lambda_{em.}$ = 445 nm) using a BioTek Synergy H1 microplate reader and comparing to a pre-determined 4-methylumbelliferyl standard curve. Enzyme activity for dilutions that produced a strictly linear profile over time were determined by calculating 4-methylumbelliferyl release (µmole $min^{-1}$) per mg of enzyme present in each reaction, averaged over relevant dilutions to give final activity of the *V. cholerae* neuraminidase stock.

**Recombinant human CD28 and human & mouse CD80.** The full-length human CD28 gene was isolated from a cDNA library from human Jurkat cells. Similarly, full-length human and mouse CD80 genes were isolated, respectively, from human Daudi cells and *ex vivo* mouse splenocytes.

Briefly, RNA extraction was performed on approximately 1 million cells (each) using the RNeasy Plus mini kit (Qiagen), according to the manufacturer's instructions. Purified total RNA was converted to a cDNA library via RT-PCR using ProtoScript II reverse transcriptase (NEB) and an 18-mer oligo-dT primer. Respective full-length CD28 and CD80 genes were amplified using Q5 DNA polymerase (NEB) and the following primer sets: Human-CD28-FWD: 5'-CTC AGG CTG CTC TTG GCT CTC AAC TTA TTC CCT TCA ATT C-3'; Human-CD28-REV: 5'-GGA GCG ATA GGC TGC GAA GTC GCG TGG TGG-3'; Human-CD80-FWD: 5'-ATG GGC CAC ACA CGG AGG CAG GGA ACA TCA C-3'; Human-CD80-REV: 5'-TAC AGG GCG TAC ACT TTC CCT TCT CAA TCT CTC ATT CC-3'; Mouse-CD80-FWD: ATG GCT TGC AAT TGT CAG TTG ATG CAG GAT ACA CC-3'; Mouse-CD80-REV: 5'-AAG GAA GAC GGT CTG TTC AGC TAA TGC TTC TTC AGG-3'. Expression clones corresponding to the respective CD28 and CD80 extracellular ectodomains (including Vset for CD28, and Vset and Cset/C2 for CD80) were amplified by PCR and subcloned via HiFi assembly (NEB) into a custom vector for mammalian cell expression encoding C-terminal eGFP and a $His_8$ tag using the following primer sets: Human-CD28-FWD: 5'-TTC CGT CCT GGC AGG ATC AAA CAA GAT TTT GGT GAA G-3'; Human-CD28-REV: 5'-CTT GAT CAG TGA TCC GGG CTT AGA AGG TCC GGG-3'; Human-CD80-FWD: 5'-GGC TTC CGT CCT GGC AGG ATC AGT TAT CCA CGT GAC CAA G-3'; Human-CD80-REV: 5'-TTT CGC GCT TGA TCA GTG ATC CTG TAT TCC AGT TGA AGG T-3'; Mouse-CD80-FWD: 5'-GGC TTC CGT CCT GGC AGG ATC AGA ACA ACT GTC CAA GTC A-3'; Mouse-CD80-REV: 5'-TTT CGC GCT TGA TCA GTG ATC CTT TTT CCC AGG TGA AGT C-3'.

For expression, purified human and mouse CD80 plasmid stocks were transfected into 293F and CHO-K1 cell cultures, respectively, using pre-complexation with PEI-MAX (Polysciences) at a 4:1 mass:mass ratio (final). 293F cells expressing human CD80 were incubated at 37°C, 8% $CO_2$, 70% humidity for 5 days before harvesting condition media for purification. Adherent CHO-K1 cells were incubated in DNA:PEI mixture for 12 hours before exchanging into serum-free media and incubation for 4 days at 37°C, 5% $CO_2$. CD80-containing media were centrifuged at low speed to remove cells and large debris, vacuum-filtered over 0.8 μm nitrocellulose membranes, and applied directly to 5mL NiNTA FF crude columns (GE), preequilibrated in 30 mM HEPES pH 7.5, 300 mM NaCl, for $Ni^{2+}$-affinity purification. Bound CD80 was eluted in a 30 mL gradient, exchanging into 30 mM HEPES pH 7.5, 300 mM NaCl, 500 mM imidazole. Protein-containing fractions were identified and purity determined via SDS-PAGE analysis. Pure fractions were pooled, concentrated to approximately 1.0 mg $mL^{-1}$ (final), aliquoted, snap frozen in liquid $N_2$, and stored at -80°C.

**Glycan microarray screening**. For monomeric proteins, recombinant protein samples (100 μg mL-1 final) were pre-complexed with anti-His mouse antibody (Thermo Fisher Scientific) and an Alexa488-linked anti-mouse IgG (Thermo Fisher Scientific) at 4:2:1 molar ratio for 15 min on ice in 100 μL Tween 20 (0.05% v/v) in PBS (PBS-T). For Fc chimeras, recombinant protein samples (100 μg mL-1 final) were pre-complexed with Alexa488- or Phycoerythrin (PE)-linked anti human IgG (Jackson) at a 2:1 molar ratio for 15 min on ice in 100 μL PBS-T. Complexes were subsequently incubated on glycan microarray slide surfaces in a humidified chamber for 60 min before washing and analysis. Following final washing, all arrays were scanned using an Innoscan 1100AL microarray scanner (Innopsys). A complete list of the glycans on the array is presented in

Extended Data Table 1. Full descriptions of the microarray experiment and datasets are presented in supplementary documentation according to the MIRAGE consortium format.

**Surface plasmon resonance (SPR)**. Direct binding of each compound to recombinant CD28 (GFP chimera) was measured by SPR using a Biacore 8K instrument (GE Healthcare) at 25 °C. Sensor chip CM5 (GE Healthcare) were used to create CD28 biosensors by standard NHS/EDC coupling. 25 µg/mL recombinant CD28 diluted in 10 mM sodium acetate (pH 5.5) was injected for 200 sec. at 5 µL/min over a single flow cell. In total, five separate CD28-coupled surfaces were generated during the course of this study with the following immobilization levels ranging from 1500-2000 resonance units (RUs). All experiments were performed in a running buffer of PBS (pH 7.4) with 0.005% (v/v) Tween 20 (PBS-T') at a flow rate of 30 µL/min. A reference flow cell was created on each sensor chip by EDC/NHS activation, followed by deactivation with ethanolamine.
To evaluate carbohydrate binding, we diluted the glycan from a 20 mM stock solution in 2X PBS-T' (7.4) to match the running buffer. Samples were injected in a dose-dependent manner (0.23 to 1 µM) over the CD28 biosensor for 30 sec followed by 120 sec of dissociation phase. The resulting sensorgrams were analyzed using Biacore 8K Evaluation Software as follows. Each sensorgram was reference subtracted by the nearest buffer blank injection. The signal just before injection stop of these corrected sensorgrams was treated as the CD28-binding response for steady-state affinity analysis and fit using four-parameter variable slope nonlinear curve fitting in GraphPad Prism8 software.

**LCMV-Clone 13.** LCMV-Clone 13 was grown, stored, and quantified according to published methods (*39*).

**Assays for functional revival of hypofunctional T cells from LCMV infected mice.** C57BL/6 mice (8–12 week old) were infected with LCMV-Clone 13 ($2x10^6$ P.F.U.) in PBS via i.v. injection. 10 days post-infection, mice were sacrificed via isoflurane overdose and subsequent cervical dislocation. Spleens were harvested and manually processed into single-cell suspensions in FACS buffer (PBS supplemented with 2% fetal bovine serum and 2 mM EDTA). Erythrocytes (RBCs) were lysed with ammonium-chloride-potassium buffer. For determination of IFN-$\gamma$ production and Granzyme B levels via intracellular staining, $4x10^6$ splenocytes were incubated in complete RPMI media (RPMI (Gibco) supplemented with 10% FBS, sodium pyruvate (Gibco), HEPES (Gibco), Pen/Strep (Gibco), β-mercaptoethanol (Gibco), and L-Glutamine (Gibco) for 5 hours in the presence or absence of 2 mg/mL GP33–41 (AnaSpec) or NP205–212 (MBL International) and/or recombinant sialidase from *V. cholerae* (55 mU). Brefeldin A (Biolegend) was added after 30 minutes of incubation. Post-incubation, cells were stained with GhostDye 510 (Tonbo) in PBS followed by surface staining in FACS buffer and subsequent fixation in 4% paraformaldehyde for 15 min.. Fixed cells were permeabilized using Perm/Wash buffer (BD) for intracellular staining.

For determination of IFN-$\gamma$ secretion, splenocytes were incubated in complete RPMI media for 3 days in the presence or absence of 2 mg/mL GP33–41 (AnaSpec) or NP205–212 (MBL International), recombinant sialidase from *V. cholerae* (55 mU), and/or 25 μg/mL anti-PD-L1 antibody (Leinco). Supernatant was collected and IFN-$\gamma$ quantified via ELISA (kit from Biolegend).

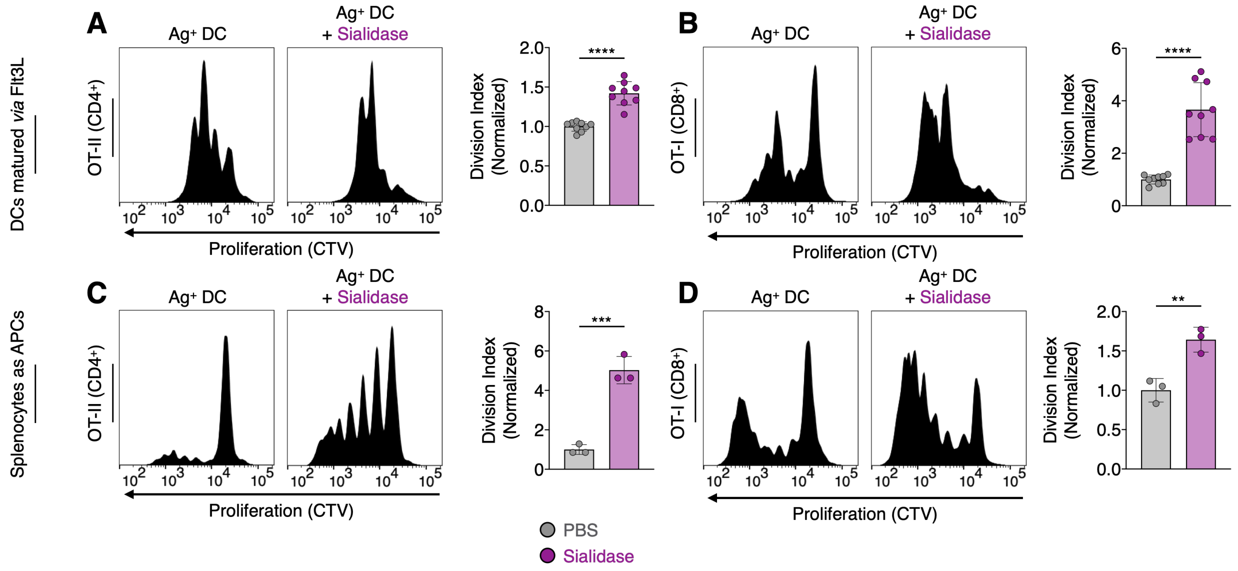


**Figure S1. Sialidase enhances antigen-dependent activation of T cells in the presence of various APCs.** T cell proliferation histograms (dilution of CTV) and quantification for co-cultures of: (**A**), CD4$^+$ OT-II cells with DCs matured from bone marrow monocytes using FMS-like tyrosine kinase 3 ligand (Flt3L); (**B**), CD8$^+$ OT-I cells with DCs matured using Flt3L (see a); (**C**), CD4$^+$ OT-II cells with bulk murine splenocytes as APCs; (**D**), CD8$^+$ OT-I cells with bulk murine splenocytes as APCs. DC (Flt3L) and splenocyte to T cell ratio was 1:2. T cell expansion was evaluated after 3 days of co-culture with APCs in the presence or absence of sialidase from *V. cholerae* (55 mU). DCs / splenocytes were preloaded with OVA (antigen, Ag) prior to co-culture with T cells. $n \geq 3$ (**A**–**D**). Mean ± s.d. (**A**–**D**). $^{**}P \leq 0.01$, $^{***}P \leq 0.001$, $^{****}P \leq 0.0001$ (**A**–**D**). One-way ANOVA followed by Tukey's multiple comparisons test (**A**–**D**). Normalized division index corresponds to T cell division index for sialidase-treated cultures divided by the division index for the corresponding PBS treated control.

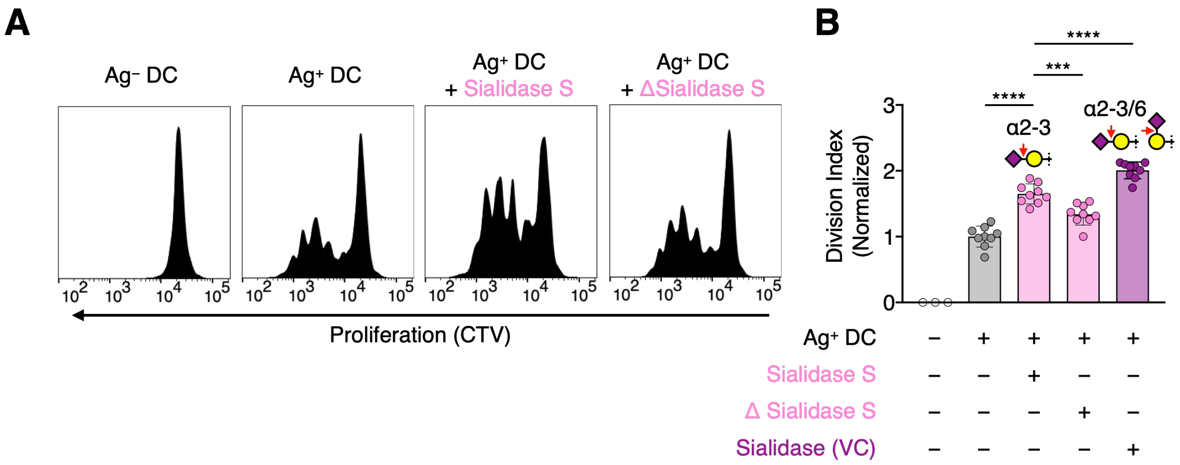


**Figure S2. The α2-3-specific sialidase from *S. pneumoniae* enhances antigen-dependent activation of CD4$^+$ T cells.** (**A**), T cell proliferation histograms (dilution of CTV) for co-cultures of CD4$^+$ OT-II cells with DCs matured from bone marrow monocytes using granulocyte-macrophage colony-stimulating factor (GM-CSF). (**B**), Quantification of data from **A**. DC (GM-CSF) and splenocyte to T cell ratio was 1:2. T cell expansion was evaluated after 3 days of co-culture with DCs in the presence or absence of sialidase from *S. pneumoniae* (Sialidase S, 0.2 mU) or *V. cholerae* (VC, 0.2 mU). DCs were preloaded with OVA (antigen, Ag) prior to co-culture with T cells. Δ = heat inactivated. $n \geq 3$ (**B**). Mean ± s.d. (**A**, **B**). ***$P \leq 0.001$, ****$P \leq 0.0001$ (**A**, **B**). One-way ANOVA followed by Tukey's multiple comparisons test (**A**, **B**). Normalized division index corresponds to T cell division index for sialidase-treated cultures divided by the division index for the corresponding PBS treated control.

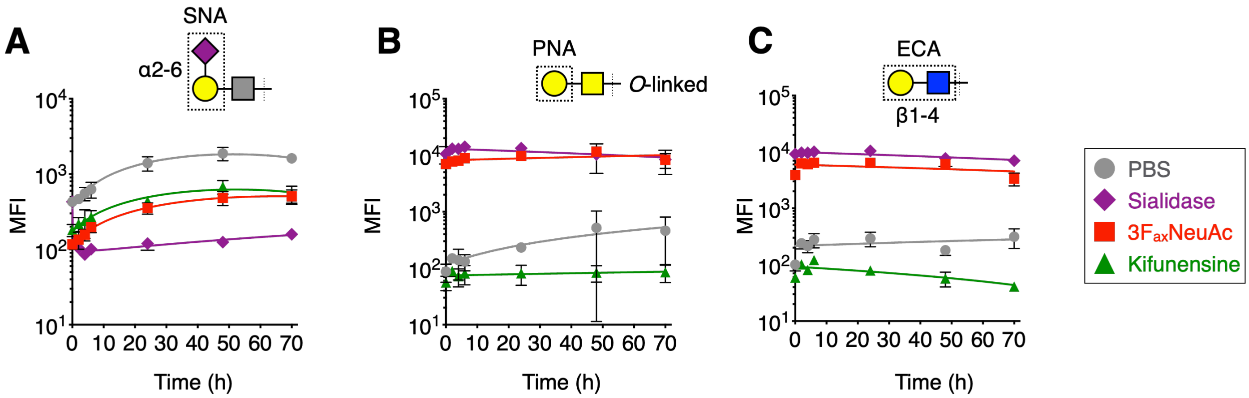


**Figure S3. Glycan dynamics on the surface of DCs treated with sialidase, $3F_{ax}$NeuAc, or kifunensine.** (**A**), DCs (GM-CSF) were treated with sialidase from *V. cholerae* (day 0), $3F_{ax}$NeuAc (day −3) , or kifunensine (day −2) and stained with fluorescent lectin from *Sambucus nigra* (SNA) at various timepoints over 72 h. (**B**), DCs were treated as in **A** and stained with fluorescent peanut agglutinin (PNA). (**C**), DCs were treated as in **A** and stained with fluorescent lectin from *Erythrina cristagalli* (ECA). Cells were analyzed *via* flow cytometry (**A**–**C**). $n = 3$ (**A**–**C**). Error bars represent mean ± s.d..

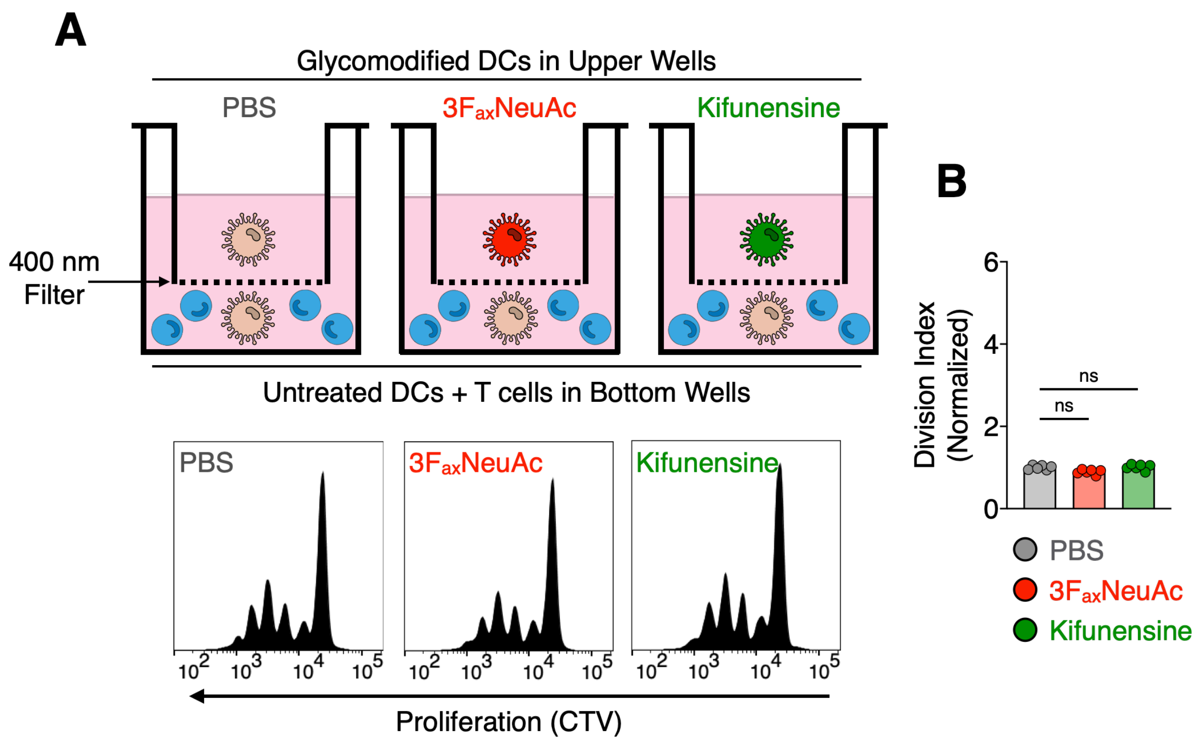


**Figure S4. Physical separation of CD4⁺ T cells from desialylated DCs abrogates enhancement in T cell activation.** (**A**), Expansion of OT-II cells in a transwell system. $3F_{ax}$NeuAc/kifunensine-treated DCs were placed in the upper well to physically separate them from normally glycosylated antigen-loaded DCs (GM-CSF) and OT-II cells in the lower well. T cell proliferation histograms at bottom. **B**, Quantification of data from **A**. Mean ± s.d. (**B**). ns = not significant (**A**, **B**). One-way ANOVA followed by Tukey's multiple comparisons test (**A**, **B**). Normalized division index corresponds to T cell division index for inhibitor-treated cultures divided by the division index for the corresponding PBS treated control. *n* = 3 (**B**).

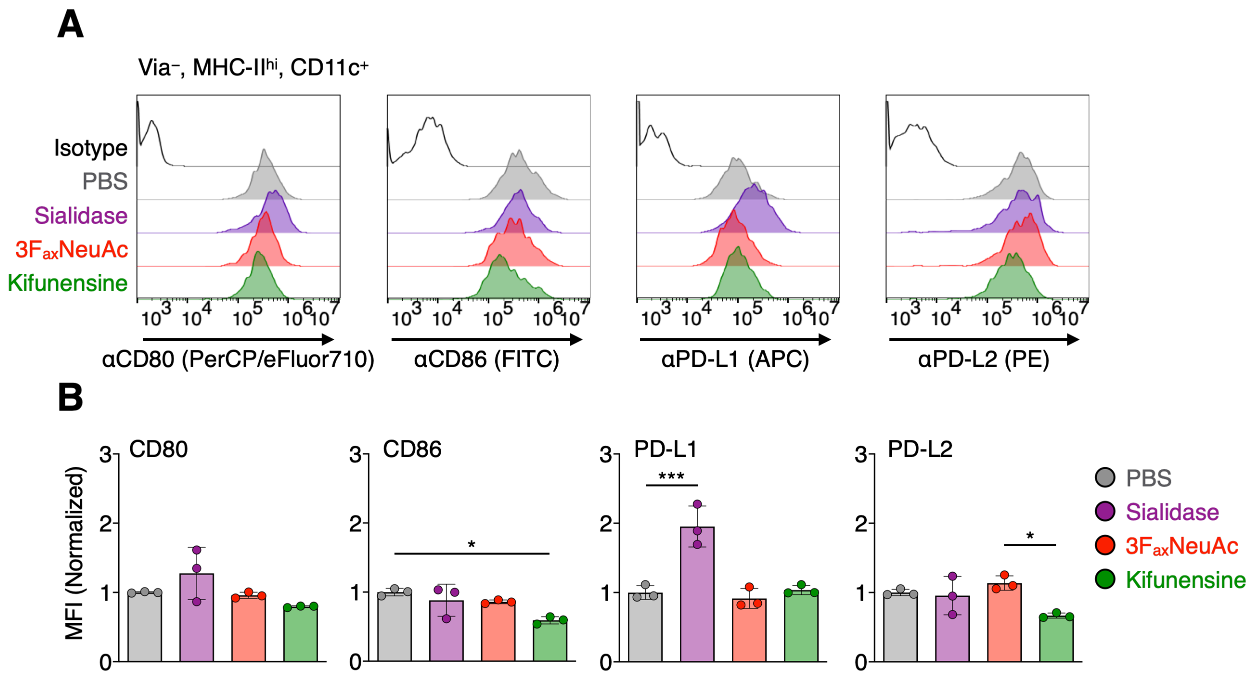


**Figure S5. Physical separation of CD4$^+$ T cells from desialylated DCs abrogates enhancement in T cell activation.** (**A**), DCs (GM-CSF) were treated with sialidase from *V. cholerae* (24 h), $3F_{ax}NeuAc$ (72 h), or kifunensine (48 h) after which protein expression was measured via flow cytometry. DCs were defined as cells expressing low viability dye (Via$^-$), high major histocompatibility complex II (MHC II$^{hi}$), and high CD11c. (**B**), Quantification of data from **A**. Peaks without statistical comparisons are not significant. Mean ± s.d. (**B**). $*P \leq 0.05$, $***P \leq 0.001$ (**B**). One-way ANOVA followed by Tukey's multiple comparisons test (**B**). Normalized MFI corresponds to division of all individual signals by the mean of the MFI for PBS treated DCs.

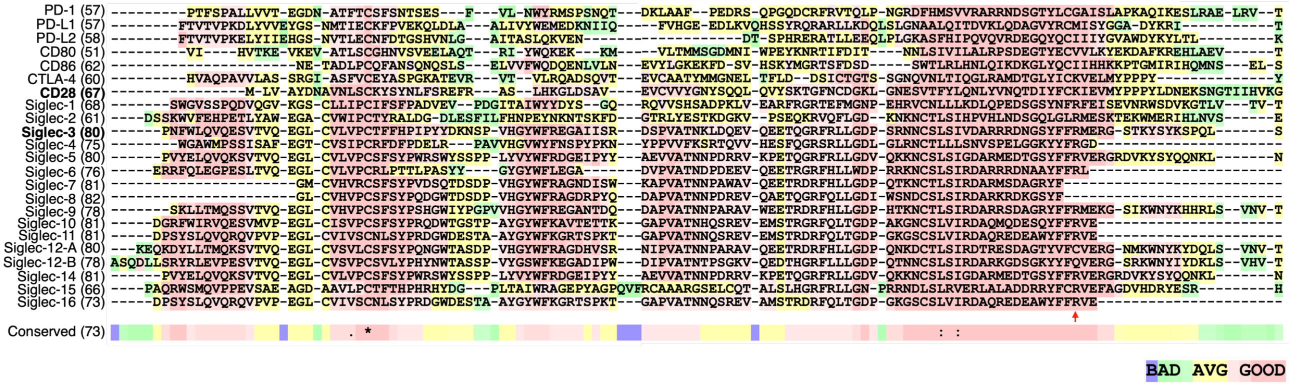


**Figure S6. Sequence alignment between receptors present at the T cell-APC immunological synapse and siglecs.** Sequences of V-set domains for all receptors were retrieved from the UniProt database. Alignments were performed using T-COFFEE Expresso tool. Alignment scores are listed in parentheses. The conserved arginine present in siglecs is highlighted by the red allow at the bottom of the alignment.

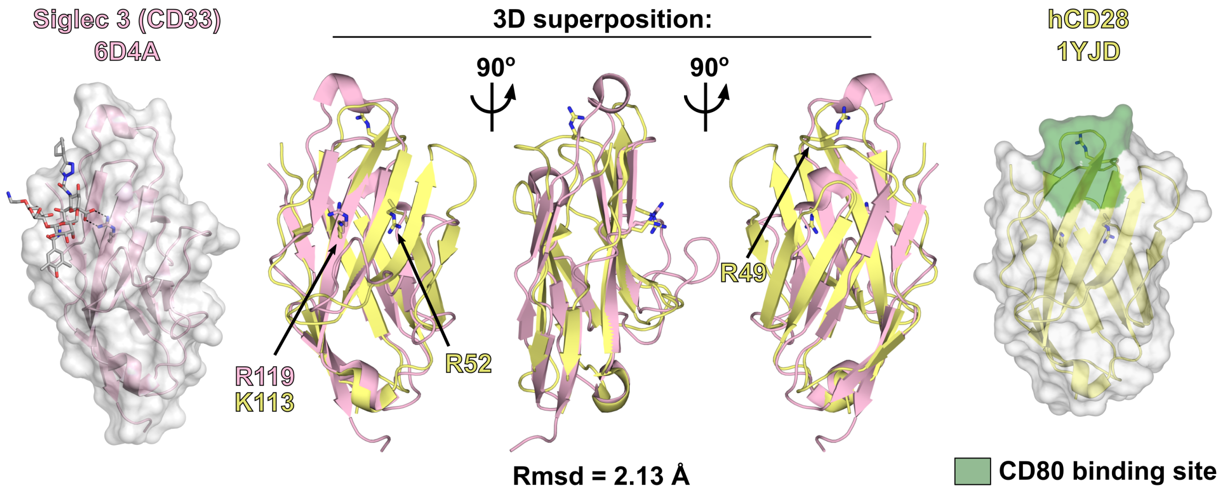


**Figure S7. Structural comparison of Siglec 3 (CD33) and human CD28.** 3D structures of the V-set domains of Siglec 3 (pink) and CD28 (light yellow) from PDB entries 6D4A and 1YJD, respectively, are shown as 3D secondary structure cartoons with partial surface overlay (far left & right panels) and as a 3D superposition of the two molecules (center panel). For Siglec 3, the binding position of a high-affinity ligand is shown as grey sticks, while the binding epitope of CD80 (MYPPY loop) is highlighted in green on the CD28 surface. The position of a key arginine (R119) required for sialic acid binding in Siglec 3 is labeled and highlighted via H-bonding interactions, while chemically/structurally similar side-chains in CD28 are shown labelled as yellow sticks. 3D superposition was performed using SSM (*40*), giving an R.m.s.d of 2.13 Å across 97 matched $C_\alpha$ positions with 10.3% sequence identity. All panels were assembled using Pymol (Schrodinger LLC).

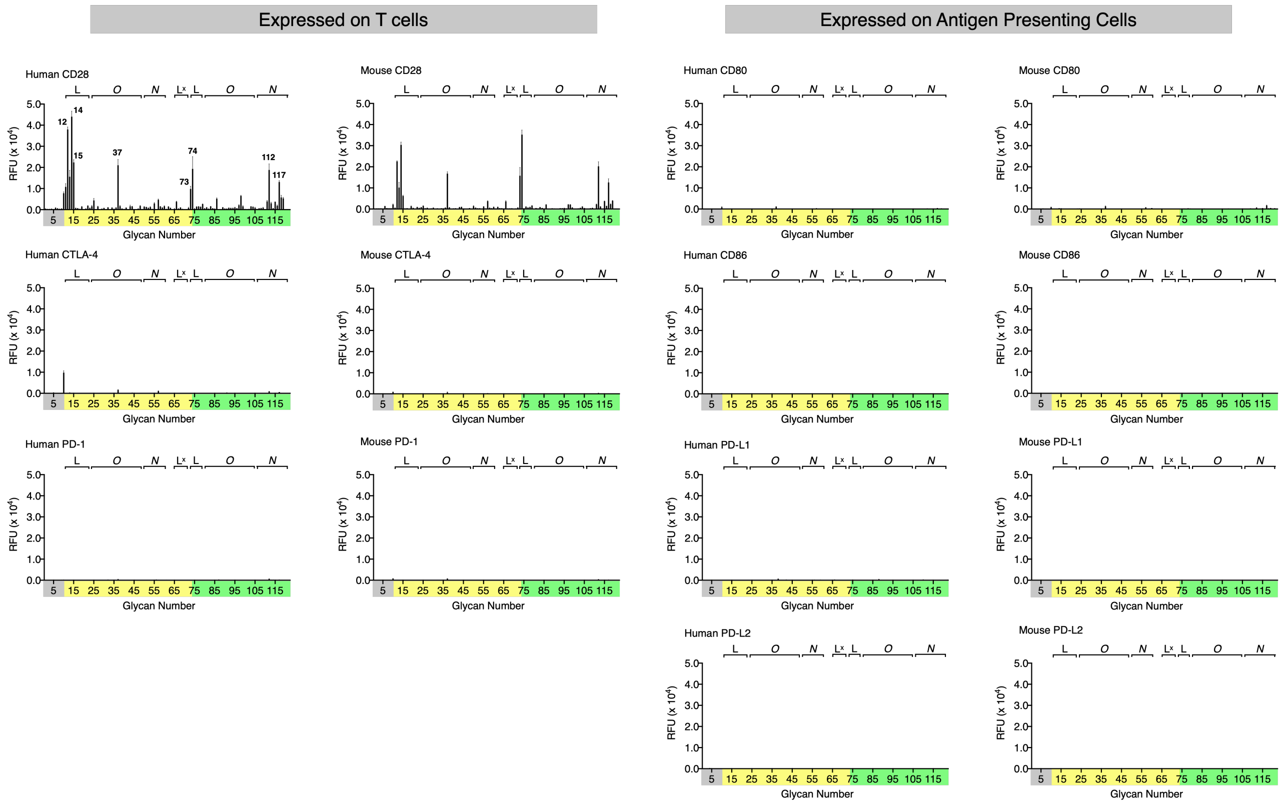


**Figure S8. Glycan microarray binding profiles of receptors present at the T cell-APC immunological synapse.** Glycan array binding data for human and mouse CD28-Fc, CTLA-4-Fc, PD-1-Fc, CD80-Fc, CD86-Fc, PD-L1-Fc, and PD-L2-Fc. Protein binding was detected using fluorescent anti-human Fc followed by microarray imaging. Compound IDs for top hits are displayed in Figure 2 in the main manuscript. $n$ = 4 for each peak. Error bars represent mean ± s.d..

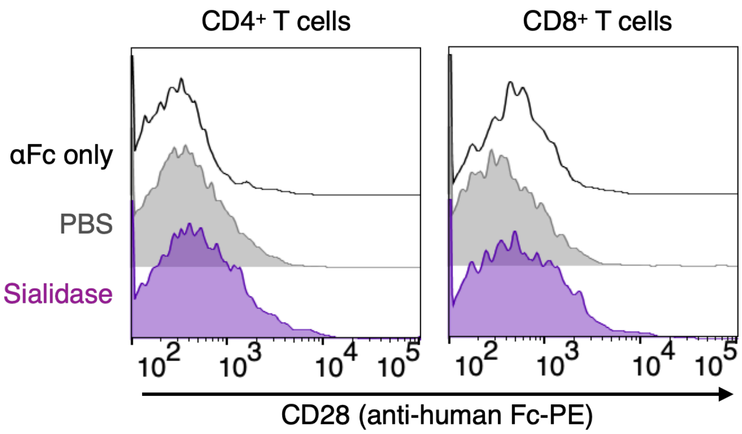


**Figure S9. CD28 does not bind to T cells regardless of sialylation status.** Murine splenocytes were treated with sialidase from *V. cholerae* (55 mU, 45 min., where appropriate) and subsequently stained with orthogonally fluorescent anti-CD3, anti-CD4, and anti-CD8 in addition to fluorescent antibodies for identifying other cell lineages. Cells were then stained with CD28-Fc (4 μg/mL) followed by fluorescent anti-human Fc. Cells were then analyzed *via* flow cytometry. T cells were defined as $CD3^{+}$ $CD4^{+}$ or $CD8^{+}$.

**Table S1. Identities of glycans on microarrays.**

| Glycan # | Common Name | Structure |
|---|---|---|
| 1 | Galβ(1-4)GlcNAcβ-ethyl-NH2 | 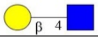 |
| 2 | Galβ(1-4)GlcNAcβ(1-3)Galβ(1-3)GalNAcα-Thr-NH2 | 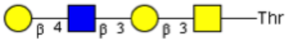  |
| 3 | Galβ(1-4)GlcNAcβ(1-6)[Galβ(1-3)]-GalNAcα-Thr-NH2 | 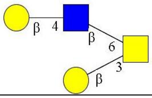 |
| 4 | Galβ(1-4)GlcNAcβ(1-3)GalNAcα-Thr-NH2 | 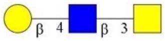 |
| 5 | Galβ(1-4)GlcNAcβ(1-3)[Galβ(1-4)GlcNAcβ(1-6)]-GalNAcα-Thr-NH2 | 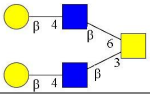 |
| 6 | Galβ(1-4)GlcNAcβ(1-6)GalNAcα-Thr-NH2 | 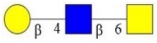 |
| 7 | Galβ(1-4)GlcNAcβ(1-2)Manα (1-3)[Galβ(1-4)GlcNAcβ(1-2)Manα(1-6)]-Manβ(1-4)GlcNAcβ(1-4)GlcNAcβ-Asn-NH2 | 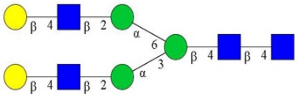 |
| 8 | Galβ(1-4)GlcNAcβ(1-2)Manα(1-3)[Galβ(1-4)GlcNAcβ(1-2)Manα(1-6)]-Manβ(1-4)GlcNAcβ(1-4)[Fucα(1-6)]-GlcNAcβ-Asn-Ser-Thr-NH2 | 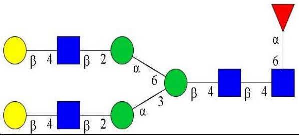 |
| 9 | Galβ(1-4)GlcNAcβ(1-2)Manα(1-3){Galβ(1-4)GlcNAcβ(1-2)[Galβ(1-4)GlcNAcβ(1-2)]-Manα(1-6)}-Manβ(1-4)GlcNAcβ(1-4)GlcNAcβ-Asn-Lys-NH2 | 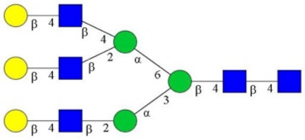 |
| 10 | Galβ(1-4)GlcNAcβ(1-2)Manα(1-3){Galβ(1-4)GlcNAcβ(1-2)[Galβ(1-4)GlcNAcβ(1-2)]-Manα(1-6)}-Manβ(1-4)GlcNAcβ(1-4)[Fucα(1-6)]-GlcNAcβ-(Lys-Val-Ala)Asn-Lys-Thr-NH2 | 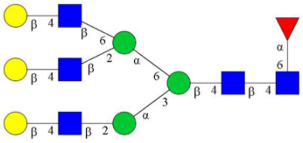 |
| 11 | NeuAcα(2-3)Galβ(1-4)6-O-sulfo-GlcNAcβ-propyl-NH2 | 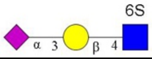  |
| 12 | NeuAcα(2-3)Galβ(1-4)[Fucα(1-3)]-6-O-sulfo-GlcNAcβ-propyl-NH2 | 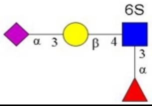  |
| 13 | NeuAcα(2-3)6-O-sulfo-Galβ(1-4)GlcNAcβ-ethyl-NH2 | 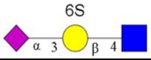  |
| 14 | NeuAcα(2-3)6-O-sulfo-Galβ(1-4)[Fucα(1-3)]-GlcNAcβ-propyl-NH2 | 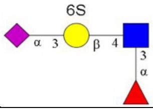  |
| 15 | NeuAcα(2-3)Galβ(1-3)6-O-sulfo-GlcNAcβ-propyl-NH2 | 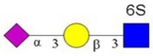  |
| 16 | NeuAcα(2-3)Galβ(1-4)Glcβ-ethyl-NH2 | 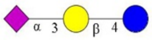 |
| 17 | NeuAcα(2-3)Galβ(1-4)GlcNAcβ-ethyl-NH2 | 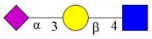 |

| Glycan # | Common Name | Structure |
|---|---|---|
| 18 | NeuAcα(2-3)Galβ(1-4)GlcNAcβ(1-3)Galβ(1-4)GlcNAcβ-ethyl-NH2 | 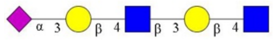 |
| 19 | NeuAcα(2-3)Galβ(1-4)GlcNAcβ(1-3)Galβ(1-4)GlcNAcβ(1-3)Galβ(1-4)GlcNAcβ-ethyl-NH2 | 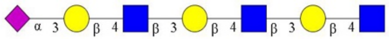 |
| 20 | NeuAcα(2-3)GalNAcβ(1-4)GlcNAcβ-ethyl-NH2 | 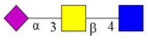 |
| 21 | NeuAcα(2-3)Galβ(1-3)GlcNAcβ-ethyl-NH2 | 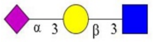 |
| 22 | NeuAcα(2-3)Galβ(1-3)GlcNAcβ(1-3)Galβ(1-4)GlcNAcβ-ethyl-NH2 | 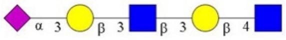 |
| 23 | NeuAcα(2-3)Galβ(1-3)GlcNAcβ(1-3)Galβ(1-3)GlcNAcβ-ethyl-NH2 | 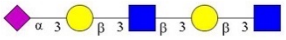 |
| 24 | NeuAcα(2-3)Galβ(1-3)GalNAcβ(1-3)Galα(1-4)Galβ(1-4)Glcβ-ethyl-NH2 | 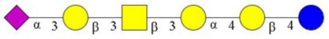 |
| 25 | NeuAcα(2-3)Galβ(1-3)GalNAcα-Thr-NH2 | 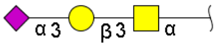 |
| 26 | NeuAcα(2-3)Galβ(1-4)GlcNAcβ(1-3)Galβ(1-3)GalNAcα-Thr-NH2 | 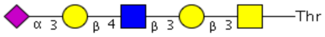 |
| 27 | NeuAcα(2-3)Galβ(1-4)GlcNAcβ(1-3)Galβ(1-4)GlcNAcβ(1-3)Galβ(1-3)GalNAcα-Thr-NH2 | 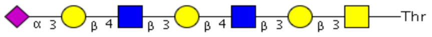 |
| 28 | NeuAcα(2-3)Galβ(1-4)GlcNAcβ(1-3)Galβ(1-4)GlcNAcβ(1-3)Galβ(1-4)GlcNAcβ(1-3)Galβ(1-3)GalNAcα-Thr-NH2 | 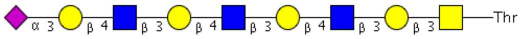 |
| 29 | NeuAcα(2-3)Galβ(1-4)GlcNAcβ(1-3)Galβ(1-4)GlcNAcβ(1-3)Galβ(1-4)GlcNAcβ(1-3)Galβ(1-4)GlcNAcβ(1-3)Galβ(1-3)GalNAcα-Thr-NH2 | 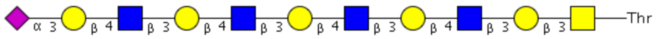 |
| 30 | NeuAcα(2-3)Galβ(1-4)GlcNAcβ(1-3)Galβ(1-4)GlcNAcβ(1-3)Galβ(1-4)GlcNAcβ(1-3)Galβ(1-4)GlcNAcβ(1-3)Galβ(1-3)GalNAcα-Thr-NH2 | 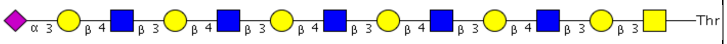 |
| 31 | NeuAcα(2-3)Galβ(1-4)GlcNAcβ(1-6)[Galβ(1-3)]-GalNAcα-Thr-NH2 | 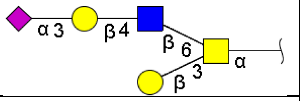 |
| 32 | NeuAcα(2-3)Galβ(1-4)GlcNAcβ(1-3)Galβ(1-4)GlcNAcβ(1-6)[Galβ(1-3)]-GalNAcα-Thr-NH2 | 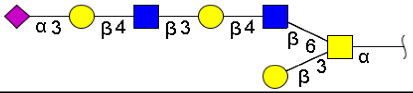 |
| 33 | NeuAcα(2-3)Galβ(1-4)GlcNAcβ(1-3)Galβ(1-4)GlcNAcβ(1-3)Galβ(1-4)GlcNAcβ(1-6)[Galβ(1-3)]-GalNAcα-Thr-NH2 | 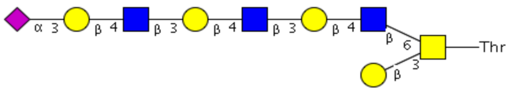 |
| 34 | NeuAcα(2-3)Galβ(1-4)GlcNAcβ(1-3)Galβ(1-4)GlcNAcβ(1-3)Galβ(1-4)GlcNAcβ(1-3)Galβ(1-4)GlcNAcβ(1-6)[Galβ(1-3)]-GalNAcα-Thr-NH2 | 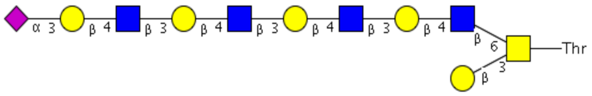 |

| Glycan # | Common Name | Structure |
|---|---|---|
| **35** | NeuAcα(2-3)Galβ(1-4)GlcNAcβ(1-3)Galβ(1-4)GlcNAcβ(1-3)Galβ(1-4)GlcNAcβ(1-3)Galβ(1-4)GlcNAcβ(1-3)Galβ(1-4)GlcNAcβ(1-6)[Galβ(1-3)]-GalNAcα-Thr-NH2 | 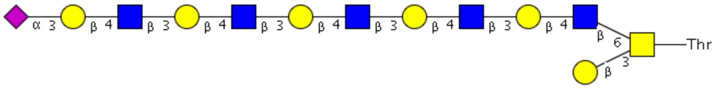 |
| **36** | NeuAcα(2-3)Galβ(1-4)GlcNAcβ(1-3)Galβ(1-4)GlcNAcβ(1-3)Galβ(1-4)GlcNAcβ(1-3)Galβ(1-4)GlcNAcβ(1-6)[NeuAcα(2-3)Galβ(1-4)GlcNAcβ(1-3)Galβ(1-4)GlcNAcβ(1-3)Galβ(1-4)GlcNAcβ(1-3)Galβ(1-3)]-GalNAcα-Thr-NH2 | 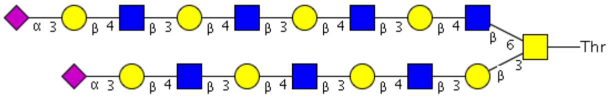 |
| **37** | NeuAcα(2-3)Galβ(1-4)GlcNAcβ(1-3)Galβ(1-4)GlcNAcβ(1-3)Galβ(1-4)GlcNAcβ(1-3)Galβ(1-4)GlcNAcβ(1-3)Galβ(1-4)GlcNAcβ(1-6)[NeuAcα(2-3)Galβ(1-4)GlcNAcβ(1-3)Galβ(1-4)GlcNAcβ(1-3)Galβ(1-4)GlcNAcβ(1-3)Galβ(1-4)GlcNAcβ(1-3)Galβ(1-3)]-GalNAcα-Thr-NH2 | 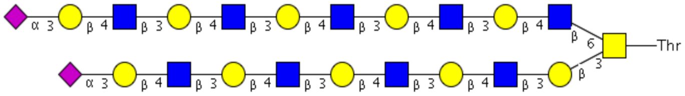 |
| **38** | NeuAcα(2-3)Galβ(1-4)GlcNAcβ(1-3)GalNAcα-Thr-NH2 | 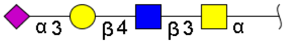 |
| **39** | NeuAcα(2-3)Galβ(1-4)GlcNAcβ(1-3)Galβ(1-4)GlcNAcβ(1-3)GalNAcα-Thr-NH2 | 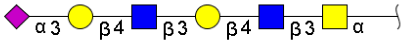 |
| **40** | NeuAcα(2-3)Galβ(1-4)GlcNAcβ(1-3)Galβ(1-4)GlcNAcβ(1-3)Galβ(1-4)GlcNAcβ(1-3)GalNAcα-Thr-NH2 | 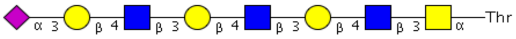 |
| **41** | NeuAcα(2-3)Galβ(1-4)GlcNAcβ(1-3)Galβ(1-4)GlcNAcβ(1-3)Galβ(1-4)GlcNAcβ(1-3)Galβ(1-4)GlcNAcβ(1-3)GalNAcα-Thr-NH2 | 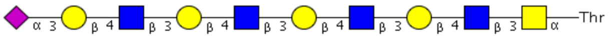 |
| **42** | NeuAcα(2-3)Galβ(1-4)GlcNAcβ(1-3)Galβ(1-4)GlcNAcβ(1-3)Galβ(1-4)GlcNAcβ(1-3)Galβ(1-4)GlcNAcβ(1-3)Galβ(1-4)GlcNAcβ(1-3)GalNAcα-Thr-NH2 | 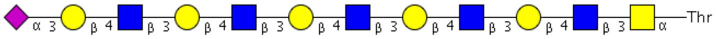 |
| **43** | NeuAcα(2-3)Galβ(1-4)GlcNAcβ(1-3)[NeuAcα(2-3)Galβ(1-4)GlcNAcβ(1-6)]-GalNAcα-Thr-NH2 | 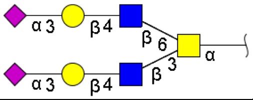 |
| **44** | NeuAcα(2-3)Galβ(1-4)GlcNAcβ(1-3)Galβ(1-4)GlcNAcβ(1-3)[NeuAcα(2-3)Galβ(1-4)GlcNAcβ(1-3)Galβ(1-4)GlcNAcβ(1-6)]-GalNAcα-Thr-NH2 | 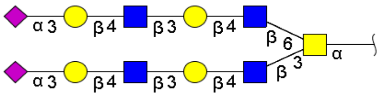 |
| **45** | NeuAcα(2-3)Galβ(1-4)GlcNAcβ(1-3)Galβ(1-4)GlcNAcβ(1-3)Galβ(1-4)GlcNAcβ(1-3)[NeuAcα(2-3)Galβ(1-4)GlcNAcβ(1-3)Galβ(1-4)GlcNAcβ(1-3)Galβ(1-4)GlcNAcβ(1-6)]-GalNAcα-Thr-NH2 | 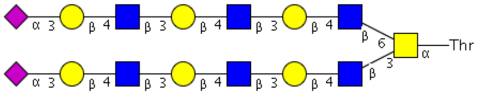 |

| Glycan # | Common Name | Structure |
| --- | --- | --- |
| **46** | NeuAcα(2-3)Galβ(1-4)GlcNAcβ(1-3)Galβ(1-4)GlcNAcβ(1-3)Galβ(1-4)GlcNAcβ(1-3)Galβ(1-4)GlcNAcβ(1-3)[NeuAcα(2-3)Galβ(1-4)GlcNAcβ(1-3)Galβ(1-4)GlcNAcβ(1-3)Galβ(1-4)GlcNAcβ(1-3)Galβ(1-4)GlcNAcβ(1-6)]-GalNAcα-Thr-NH2 | 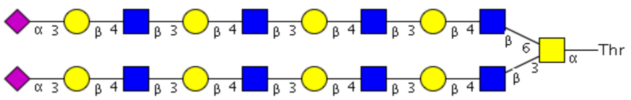  |
| **47** | NeuAcα(2-3)Galβ(1-4)GlcNAcβ(1-3)Galβ(1-4)GlcNAcβ(1-3)Galβ(1-4)GlcNAcβ(1-3)Galβ(1-4)GlcNAcβ(1-3)Galβ(1-4)GlcNAcβ(1-3)[NeuAcα(2-3)Galβ(1-4)GlcNAcβ(1-3)Galβ(1-4)GlcNAcβ(1-3)Galβ(1-4)GlcNAcβ(1-3)Galβ(1-4)GlcNAcβ(1-3)Galβ(1-4)GlcNAcβ(1-6)]-GalNAcα-Thr-NH2 | 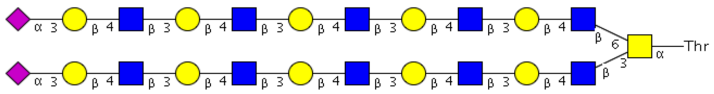  |
| **48** | NeuAcα(2-3)Galβ(1-4)GlcNAcβ(1-3)Galβ(1-4)GlcNAcβ(1-3)Galβ(1-4)GlcNAcβ(1-3)Galβ(1-4)GlcNAcβ(1-6)GalNAcα-Thr-NH2 | 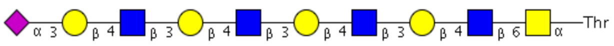  |
| **49** | NeuAcα(2-3)Galβ(1-4)GlcNAcβ(1-3)Galβ(1-4)GlcNAcβ(1-3)Galβ(1-4)GlcNAcβ(1-3)Galβ(1-4)GlcNAcβ(1-3)Galβ(1-4)GlcNAcβ(1-6)GalNAcα-Thr-NH2 | 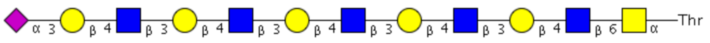  |
| **50** | NeuAcα(2-3)Galβ(1-3)GlcNAcβ(1-3)Galβ(1-4)GlcNAcβ(1-6)[NeuAcα(2-3)Galβ(1-3)GlcNAcβ(1-3)] Galβ(1-4)GlcNAcβ-ethyl-NH2 | 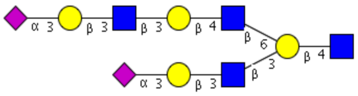 |
| **51** | NeuAcα(2-3)Galβ(1-4)GlcNAcβ(1-2)Manα(1-3)[NeuAcα(2-3)Galβ(1-4)GlcNAcβ(1-2)Manα(1-6)]-Manβ(1-4)GlcNAcβ(1-4)GlcNAcβ-Asn-NH2 | 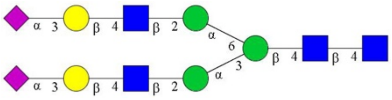 |
| **52** | NeuAcα(2-3)Galβ(1-4)GlcNAcβ(1-3)Galβ(1-4)GlcNAcβ(1-2)Manα(1-3)[NeuAcα(2-3)Galβ(1-4)GlcNAcβ(1-3)Galβ(1-4)GlcNAcβ(1-2)Manα(1-6)]-Manβ(1-4)GlcNAcβ(1-4)GlcNAcβ-Asn-NH2 | 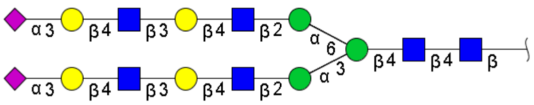 |
| **53** | NeuAcα(2-3)Galβ(1-4)GlcNAcβ(1-3)Galβ(1-4)GlcNAcβ(1-3)Galβ(1-4)GlcNAcβ(1-2)Manα(1-3)[NeuAcα(2-3)Galβ(1-4)GlcNAcβ(1-3)Galβ(1-4)GlcNAcβ(1-3)Galβ(1-4)GlcNAcβ(1-2)Manα(1-6)]-Manβ(1-4)GlcNAcβ(1-4)GlcNAcβ-Asn-NH2 | 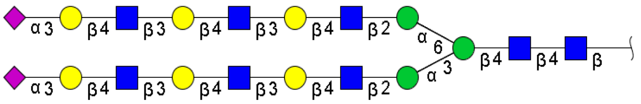 |
| **54** | NeuAcα(2-3)Galβ(1-4)GlcNAcβ(1-3)Galβ(1-4)GlcNAcβ(1-2)Manα(1-3)[NeuAcα(2-3)Galβ(1-4)GlcNAcβ(1-3)Galβ(1-4)GlcNAcβ(1-2)Manα(1-6)]-Manβ(1-4)GlcNAcβ(1-4)GlcNAcβ-(Lys-Val-Ala)Asn-Lys-Thr-NH2 | 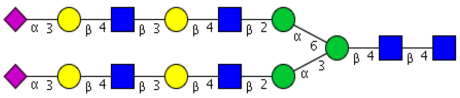 |

| Glycan # | Common Name | Structure |
|---|---|---|
| **55** | NeuAcα(2-3)Galβ(1-4)GlcNAcβ(1-3)Galβ(1-4)GlcNAcβ(1-3)Galβ(1-4)GlcNAcβ(1-2)Manα(1-3)[NeuAcα(2-3)Galβ(1-4)GlcNAcβ(1-3)Galβ(1-4)GlcNAcβ(1-3)Galβ(1-4)GlcNAcβ(1-2)Manα(1-6)]-Manβ(1-4)GlcNAcβ(1-4)GlcNAcβ-(Lys-Val-Ala)Asn-Lys-Thr-NH2 | 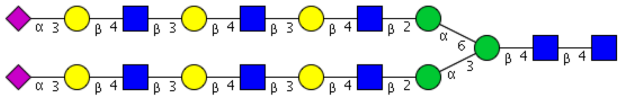 |
| **56** | NeuAcα(2-3)Galβ(1-4)GlcNAcβ(1-3)Galβ(1-4)GlcNAcβ(1-3)Galβ(1-4)GlcNAcβ(1-3)Galβ(1-4)GlcNAcβ(1-2)Manα(1-3)[NeuAcα(2-3)Galβ(1-4)GlcNAcβ(1-3)Galβ(1-4)GlcNAcβ(1-3)Galβ(1-4)GlcNAcβ(1-3)Galβ(1-4)GlcNAcβ(1-2)Manα(1-6)]-Manβ(1-4)GlcNAcβ(1-4)GlcNAcβ-(Lys-Val-Ala)Asn-Lys-Thr-NH2 | 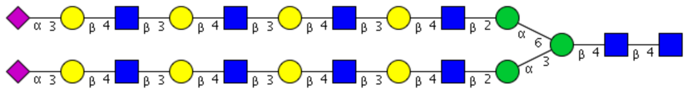 |
| **57** | NeuAcα(2-3)Galβ(1-4)GlcNAcβ(1-3)Galβ(1-4)GlcNAcβ(1-3)Galβ(1-4)GlcNAcβ(1-2)Manα(1-3)[NeuAcα(2-3)Galβ(1-4)GlcNAcβ(1-3)Galβ(1-4)GlcNAcβ(1-3)Galβ(1-4)GlcNAcβ(1-2)Manα(1-6)]-Manβ(1-4)GlcNAcβ(1-4)[Fucα(1-6)]-GlcNAcβ-(Lys-Val-Ala)Asn-Lys-Thr-NH2 | 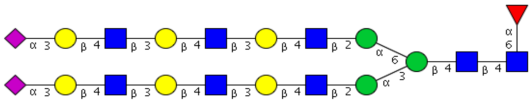 |
| **58** | NeuAcα(2-3)Galβ(1-4)GlcNAcβ(1-3)Galβ(1-4)GlcNAcβ(1-3)Galβ(1-4)GlcNAcβ(1-3)Galβ(1-4)GlcNAcβ(1-2)Manα(1-3)[NeuAcα(2-3)Galβ(1-4)GlcNAcβ(1-3)Galβ(1-4)GlcNAcβ(1-3)Galβ(1-4)GlcNAcβ(1-3)Galβ(1-4)GlcNAcβ(1-2)Manα(1-6)]-Manβ(1-4)GlcNAcβ(1-4)[Fucα(1-6)]-GlcNAcβ-(Lys-Val-Ala)Asn-Lys-Thr-NH2 | 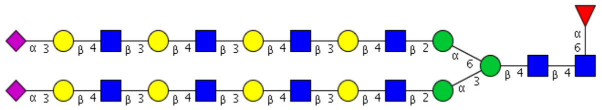 |
| **59** | NeuAcα(2-3)Galβ(1-4)GlcNAcβ(1-3)Galβ(1-4)GlcNAcβ(1-2)Manα(1-3){NeuAcα(2-3)Galβ(1-4)GlcNAcβ(1-3)Galβ(1-4)GlcNAcβ(1-2)[NeuAcα(2-3)Galβ(1-4)GlcNAcβ(1-3)Galβ(1-4)GlcNAcβ(1-6)Manα(1-6)]}-Manβ(1-4)GlcNAcβ(1-4)GlcNAcβ-(Lys-Val-Ala)Asn-Lys-Thr-NH2 | 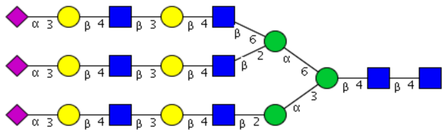 |
| **60** | NeuAcα(2-3)Galβ(1-4)GlcNAcβ(1-3)Galβ(1-4)GlcNAcβ(1-2)Manα(1-3){NeuAcα(2-3)Galβ(1-4)GlcNAcβ(1-3)Galβ(1-4)GlcNAcβ(1-2)[NeuAcα(2-3)Galβ(1-4)GlcNAcβ(1-3)Galβ(1-4)GlcNAcβ(1-6)Manα(1-6)]}-Manβ(1-4)GlcNAcβ(1-4)[Fucα(1-6)]-GlcNAcβ-(Lys-Val-Ala)Asn-Lys-Thr-NH2 | 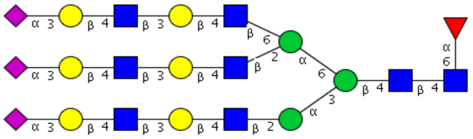 |

| Glycan # | Common Name | Structure |
|---|---|---|
| **61** | Gn/3'SLN/3'SLN-TriN | 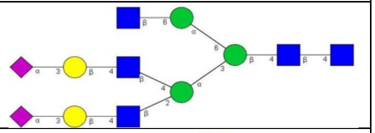 |
| **62** | NeuAcα(2-3)[GalNAcβ(1-4)]-Galβ(1-4)GlcNAcβ-ethyl-NH2 | 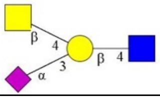 |
| **63** | NeuAcα(2-3)[GalNAcβ(1-4)]-Galβ(1-4)Glcβ-ethyl-NH2 | 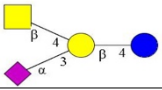 |
| **64** | Galβ(1-3)GalNAcβ(1-4)[NeuAcα(2-3)]-Galβ(1-4)Glcβ-ethyl-NH2 | 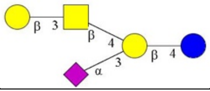 |
| **65** | NeuAcα(2-3)Galβ(1-4)[Fucα(1-3)]-GlcNAcβ-propyl-NH2 | 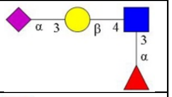 |
| **66** | NeuAcα(2-3)Galβ(1-3)[Fucα(1-4)]-GlcNAcβ(1-3)Galβ(1-4)[Fucα(1-3)]-GlcNAcβ-ethyl-NH2 | 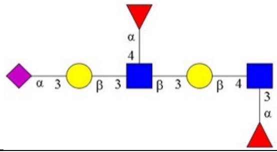 |
| **67** | NeuAcα(2-3)Galβ(1-4)[Fucα(1-3)]-GlcNAcβ(1-3)Galβ(1-4)[Fucα(1-3)]-GlcNAcβ-ethyl-NH2 | 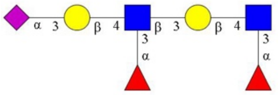 |
| **68** | NeuAcα(2-3)Galβ(1-4)[Fucα(1-3)]-GlcNAcβ(1-3)Galβ(1-4)[Fucα(1-3)]-GlcNAcβ(1-3)Galβ(1-4)[Fucα(1-3)]-GlcNAcβ-ethyl-NH2 | 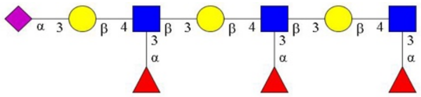 |
| **69** | NeuAcα(2-3)Galβ(1-4)[Fucα(1-3)]-GlcNAcβ(1-3)Galβ(1-4)[Fucα(1-3)]-GlcNAcβ(1-3)Galβ(1-4)[Fucα(1-3)]-GlcNAcβ(1-3)Galβ(1-3)GalNAcα-Thr-NH2 | 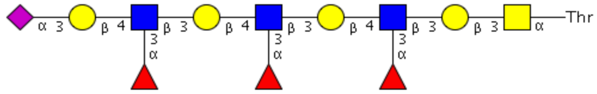 |
| **70** | NeuAcα(2-3)Galβ(1-4)[Fucα(1-3)]-GlcNAcβ(1-3)Galβ(1-4)[Fucα(1-3)]-GlcNAcβ(1-3)Galβ(1-4)[Fucα(1-3)]-GlcNAcβ(1-3)GalNAcα-Thr-NH2 | 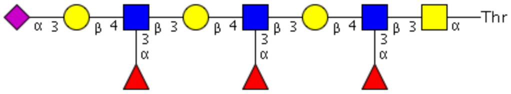 |
| **71** | NeuAcα(2-3)Galβ(1-4)[Fucα(1-3)]-GlcNAcβ(1-3)Galβ(1-4)[Fucα(1-3)]-GlcNAcβ(1-3)Galβ(1-4)[Fucα(1-3)]-GlcNAcβ(1-3)[NeuAcα(2-3)Galβ(1-4)[Fucα(1-3)]-GlcNAcβ(1-3)Galβ(1-4)[Fucα(1-3)]-GlcNAcβ(1-3)Galβ(1-4)[Fucα(1-3)]-GlcNAcβ(1-6)]-GalNAcα-Thr-NH2 | 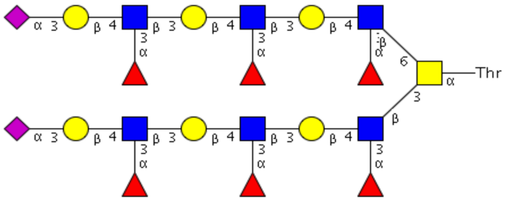 |
| **72** | NeuAcα(2-3)Galβ(1-4)[Fucα(1-3)]-GlcNAcβ(1-2)Manα(1-3)[NeuAcα(2-3)Galβ(1-4)[Fucα(1-3)]-GlcNAcβ(1-2)Manα(1-6)]-Manβ(1-4)GlcNAcβ(1-4)GlcNAcβ-(Lys-Val-Ala)Asn-(Lys-Thr)NH2 | 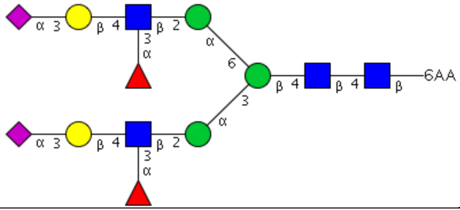 |

| Glycan # | Common Name | Structure |
|---|---|---|
| 73 | NeuAcα(2-6)Galb(1-4)(6S)GlcNacb-ethyl-NH2 | 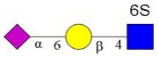 |
| 74 | NeuAcα(2-6)Galβ(1-4)6-O-sulfo-GlcNAcβ-propyl-NH2 | 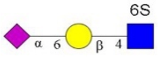 |
| 75 | NeuAcα(2-6)Galβ(1-4)Glcβ-ethyl-NH2 | 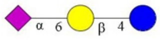 |
| 76 | NeuAcα(2-6)Galβ(1-4)GlcNAcβ-ethyl-NH2 | 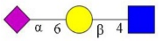 |
| 77 | NeuAcα(2-6)Galβ(1-4)GlcNAcβ(1-3)Galβ(1-4)GlcNAcβ-ethyl-NH2 | 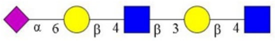 |
| 78 | NeuAcα(2-6)Galβ(1-4)GlcNAcβ(1-3)Galβ(1-4)GlcNAcβ(1-3)Galβ(1-4)GlcNAcβ-ethyl-NH2 | 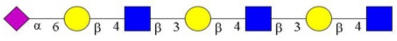 |
| 79 | NeuAcα(2-6)GalNAcβ(1-4)GlcNAcβ-ethyl-NH2 | 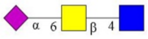 |
| 80 | NeuAcα(2-6)Galβ(1-4)GlcNAcβ(1-3)Galβ(1-3)GalNAcα-Thr-NH2 | 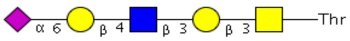 |
| 81 | NeuAcα(2-6)Galβ(1-4)GlcNAcβ(1-3)Galβ(1-4)GlcNAcβ(1-3)Galβ(1-3)GalNAcα-Thr-NH2 | 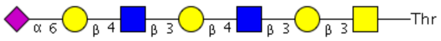 |
| 82 | NeuAcα(2-6)Galβ(1-4)GlcNAcβ(1-3)Galβ(1-4)GlcNAcβ(1-3)Galβ(1-4)GlcNAcβ(1-3)Galβ(1-3)GalNAcα-Thr-NH2 | 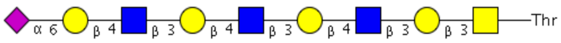 |
| 83 | NeuAcα(2-6)Galβ(1-4)GlcNAcβ(1-3)Galβ(1-4)GlcNAcβ(1-3)Galβ(1-4)GlcNAcβ(1-3)Galβ(1-4)GlcNAcβ(1-3)Galβ(1-3)GalNAcα-Thr-NH2 | 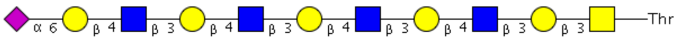 |
| 84 | NeuAcα(2-6)Galβ(1-4)GlcNAcβ(1-3)Galβ(1-4)GlcNAcβ(1-3)Galβ(1-4)GlcNAcβ(1-3)Galβ(1-4)GlcNAcβ(1-3)Galβ(1-4)GlcNAcβ(1-3)Galβ(1-3)GalNAcα-Thr-NH2 | 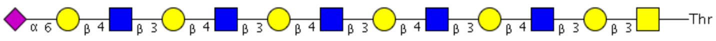 |
| 85 | NeuAcα(2-6)Galβ(1-4)GlcNAcβ(1-6)[Galβ(1-3)]-GalNAcα-Thr-NH2 | 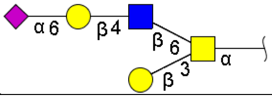 |
| 86 | NeuAcα(2-6)Galβ(1-4)GlcNAcβ(1-3)Galβ(1-4)GlcNAcβ(1-6)[Galβ(1-3)]-GalNAcα-Thr-NH2 | 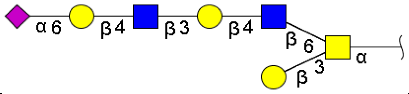 |
| 87 | NeuAcα(2-6)Galβ(1-4)GlcNAcβ(1-3)Galβ(1-4)GlcNAcβ(1-3)Galβ(1-4)GlcNAcβ(1-6)[Galβ(1-3)]-GalNAcα-Thr-NH2 | 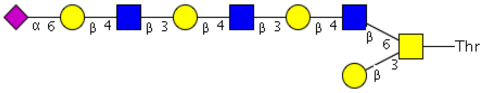 |
| 88 | NeuAcα(2-6)Galβ(1-4)GlcNAcβ(1-3)Galβ(1-4)GlcNAcβ(1-3)Galβ(1-4)GlcNAcβ(1-3)Galβ(1-4)GlcNAcβ(1-6)[Galβ(1-3)]-GalNAcα-Thr-NH2 | 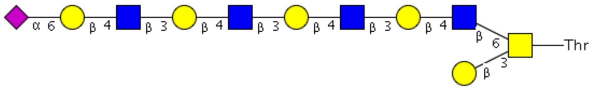 |
| 89 | NeuAcα(2-6)Galβ(1-4)GlcNAcβ(1-3)Galβ(1-4)GlcNAcβ(1-3)Galβ(1-4)GlcNAcβ(1-3)Galβ(1-4)GlcNAcβ(1-3)Galβ(1-4)GlcNAcβ(1-6)[Galβ(1-3)]-GalNAcα-Thr-NH2 | 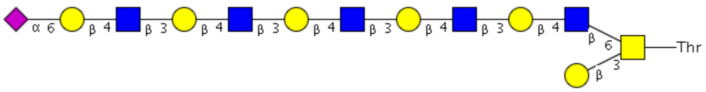 |

| Glycan # | Common Name | Structure |
|---|---|---|
| **90** | NeuAcα(2-6)Galβ(1-4)GlcNAcβ(1-3)Galβ(1-4)GlcNAcβ(1-3)Galβ(1-4)GlcNAcβ(1-3)Galβ(1-4)GlcNAcβ(1-6)[NeuAcα(2-6)Galβ(1-4)GlcNAcβ(1-3)Galβ(1-4)GlcNAcβ(1-3)Galβ(1-4)GlcNAcβ(1-3)Galβ(1-3)]-GalNAcα-Thr-NH2 | 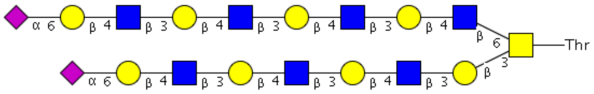 |
| **91** | NeuAcα(2-6)Galβ(1-4)GlcNAcβ(1-3)Galβ(1-4)GlcNAcβ(1-3)Galβ(1-4)GlcNAcβ(1-3)Galβ(1-4)GlcNAcβ(1-3)Galβ(1-4)GlcNAcβ(1-6)[NeuAcα(2-6)Galβ(1-4)GlcNAcβ(1-3)Galβ(1-4)GlcNAcβ(1-3)Galβ(1-4)GlcNAcβ(1-3)Galβ(1-4)GlcNAcβ(1-3)Galβ(1-3)]-GalNAcα-Thr-NH2 | 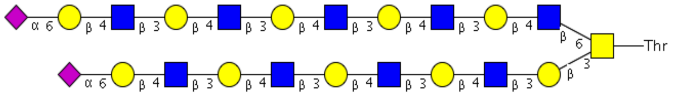 |
| **92** | NeuAcα(2-6)Galβ(1-4)GlcNAcβ(1-3)GalNAcα-Thr-NH2 | 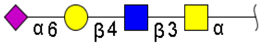 |
| **93** | NeuAcα(2-6)Galβ(1-4)GlcNAcβ(1-3)Galβ(1-4)GlcNAcβ(1-3)GalNAcα-Thr-NH2 | 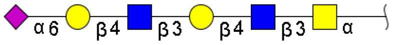 |
| **94** | NeuAcα(2-6)Galβ(1-4)GlcNAcβ(1-3)Galβ(1-4)GlcNAcβ(1-3)Galβ(1-4)GlcNAcβ(1-3)GalNAcα-Thr-NH2 | 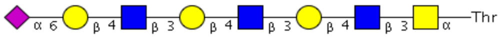 |
| **95** | NeuAcα(2-6)Galβ(1-4)GlcNAcβ(1-3)Galβ(1-4)GlcNAcβ(1-3)Galβ(1-4)GlcNAcβ(1-3)Galβ(1-4)GlcNAcβ(1-3)GalNAcα-Thr-NH2 | 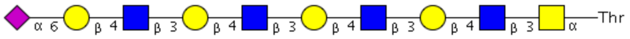 |
| **96** | NeuAcα(2-6)Galβ(1-4)GlcNAcβ(1-3)Galβ(1-4)GlcNAcβ(1-3)Galβ(1-4)GlcNAcβ(1-3)Galβ(1-4)GlcNAcβ(1-3)Galβ(1-4)GlcNAcβ(1-3)GalNAcα-Thr-NH2 | 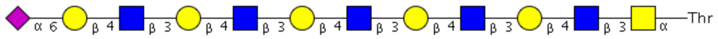 |
| **97** | NeuAcα(2-6)Galβ(1-4)GlcNAcβ(1-3)[NeuAcα(2-6)Galβ(1-4)GlcNAcβ(1-6)]-GalNAcα-Thr-NH2 | 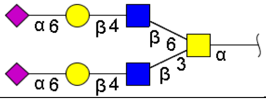 |
| **98** | NeuAcα(2-6)Galβ(1-4)GlcNAcβ(1-3)Galβ(1-4)GlcNAcβ(1-3)[NeuAcα(2-6)Galβ(1-4)GlcNAcβ(1-3)Galβ(1-4)GlcNAcβ(1-6)]-GalNAcα-Thr-NH2 | 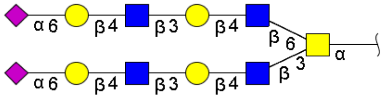 |
| **99** | NeuAcα(2-6)Galβ(1-4)GlcNAcβ(1-3)Galβ(1-4)GlcNAcβ(1-3)Galβ(1-4)GlcNAcβ(1-3)[NeuAcα(2-6)Galβ(1-4)GlcNAcβ(1-3)Galβ(1-4)GlcNAcβ(1-3)Galβ(1-4)GlcNAcβ(1-6)]-GalNAcα-Thr-NH2 | 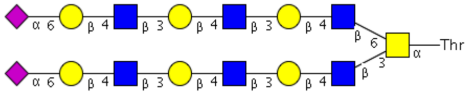 |
| **100** | NeuAcα(2-6)Galβ(1-4)GlcNAcβ(1-3)Galβ(1-4)GlcNAcβ(1-3)Galβ(1-4)GlcNAcβ(1-3)Galβ(1-4)GlcNAcβ(1-3)[NeuAcα(2-6)Galβ(1-4)GlcNAcβ(1-3)Galβ(1-4)GlcNAcβ(1-3)Galβ(1-4)GlcNAcβ(1-3)Galβ(1-4)GlcNAcβ(1-6)]-GalNAcα-Thr-NH2 | 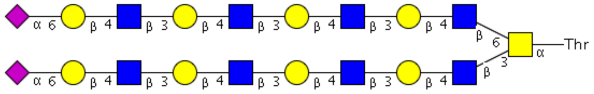 |

| Glycan # | Common Name | Structure |
|---|---|---|
| **101** | NeuAcα(2-6)Galβ(1-4)GlcNAcβ(1-3)Galβ(1-4)GlcNAcβ(1-3)Galβ(1-4)GlcNAcβ(1-3)Galβ(1-4)GlcNAcβ(1-3)Galβ(1-4)GlcNAcβ(1-3)[NeuAcα(2-6)Galβ(1-4)GlcNAcβ(1-3)Galβ(1-4)GlcNAcβ(1-3)Galβ(1-4)GlcNAcβ(1-3)Galβ(1-4)GlcNAcβ(1-3)Galβ(1-4)GlcNAcβ(1-6)]-GalNAcα-Thr-NH2 | 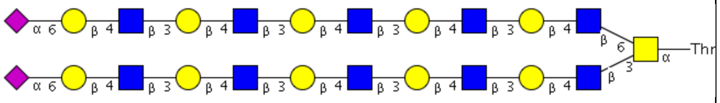 |
| **102** | NeuAcα(2-6)Galβ(1-4)GlcNAcβ(1-3)Galβ(1-4)GlcNAcβ(1-3)Galβ(1-4)GlcNAcβ(1-3)Galβ(1-4)GlcNAcβ(1-6)GalNAcα-Thr-NH2 | 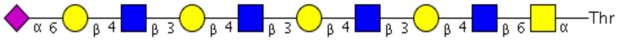 |
| **103** | NeuAcα(2-6)Galβ(1-4)GlcNAcβ(1-3)Galβ(1-4)GlcNAcβ(1-3)Galβ(1-4)GlcNAcβ(1-3)Galβ(1-4)GlcNAcβ(1-3)Galβ(1-4)GlcNAcβ(1-6)GalNAcα-Thr-NH2 | 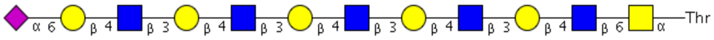 |
| **104** | NeuAcα(2-6)Galβ(1-4)GlcNAcβ(1-3)Galβ(1-4)GlcNAcβ(1-3)Galβ(1-4)GlcNAcβ(1-6)[NeuAcα(2-6)Galβ(1-4)GlcNAcβ(1-3)Galβ(1-4)GlcNAcβ(1-3)] Galβ(1-4)GlcNAcβ-ethyl-NH2 | 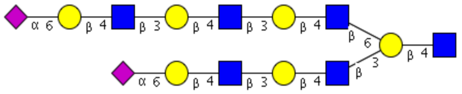 |
| **105** | NeuAcα(2-6)Galβ(1-3)GlcNAcβ(1-3)Galβ(1-4)GlcNAcβ(1-6)[NeuAcα(2-6)Galβ(1-3)GlcNAcβ(1-3)] Galβ(1-4)GlcNAcβ-ethyl-NH2 | 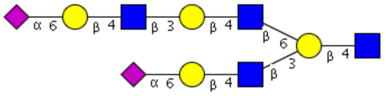 |
| **106** | Galβ(1-4)GlcNAcβ(1-2)Manα(1-3)[NeuAcα(2-6)Galβ(1-4)GlcNAcβ(1-2)Manα(1-6)]-Manβ(1-4)GlcNAcβ(1-4)GlcNAcβ-Asn-NH2 | 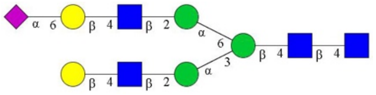 |
| **107** | NeuAcα(2-6)Galβ(1-4)GlcNAcβ(1-2)Manα(1-3)[Galβ(1-4)GlcNAcβ(1-2)Manα(1-6)]-Manβ(1-4)GlcNAcβ(1-4)GlcNAcβ-Asn-NH2 | 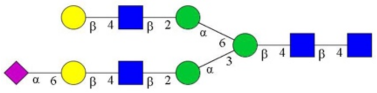 |
| **108** | GlcNAcβ(1-2)Manα(1-3)[NeuAcα(2-6)Galβ(1-4)GlcNAcβ(1-2)Manα(1-6)]-Manβ(1-4)GlcNAcβ(1-4)GlcNAcβ-Asn-NH2 | 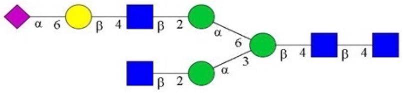 |
| **109** | NeuAcα(2-6)Galβ(1-4)GlcNAcβ(1-2)Manα(1-3)[NeuAcα(2-6)Galβ(1-4)GlcNAcβ(1-2)Manα(1-6)]-Manβ(1-4)GlcNAcβ(1-4)GlcNAcβ-Asn-NH2 | 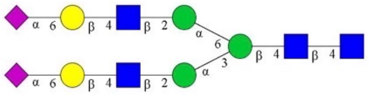 |
| **110** | NeuAcα(2-6)Galβ(1-4)GlcNAcβ(1-3)Galβ(1-4)GlcNAcβ(1-2)Manα(1-3)[NeuAcα(2-6)Galβ(1-4)GlcNAcβ(1-3)Galβ(1-4)GlcNAcβ(1-2)Manα(1-6)]-Manβ(1-4)GlcNAcβ(1-4)GlcNAcβ-Asn-NH2 | 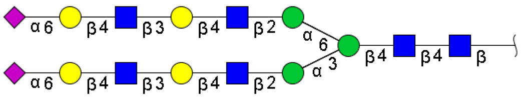 |

| Glycan # | Common Name | Structure |
|---|---|---|
| **111** | NeuAcα(2-6)Galβ(1-4)GlcNAcβ(1-3)Galβ(1-4)GlcNAcβ(1-2)Manα(1-3)[NeuAcα(2-6)Galβ(1-4)GlcNAcβ(1-3)Galβ(1-4)GlcNAcβ(1-2)Manα(1-6)]-Manβ(1-4)GlcNAcβ(1-4)GlcNAcβ-(Lys-Val-Ala)Asn-Lys-Thr-NH2 | 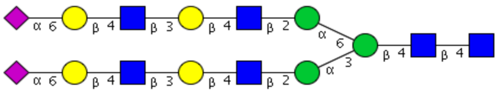 |
| **112** | NeuAcα(2-6)Galβ(1-4)GlcNAcβ(1-3)Galβ(1-4)GlcNAcβ(1-3)Galβ(1-4)GlcNAcβ(1-2)Manα(1-3)[NeuAcα(2-6)Galβ(1-4)GlcNAcβ(1-3)Galβ(1-4)GlcNAcβ(1-3)Galβ(1-4)GlcNAcβ(1-2)Manα(1-6)]-Manβ(1-4)GlcNAcβ(1-4)GlcNAcβ-Asn-NH2 | 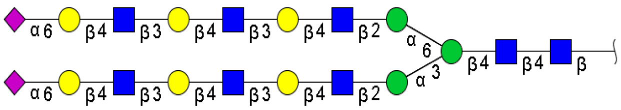 |
| **113** | NeuAcα(2-6)Galβ(1-4)GlcNAcβ(1-3)Galβ(1-4)GlcNAcβ(1-3)Galβ(1-4)GlcNAcβ(1-2)Manα(1-3)[NeuAcα(2-6)Galβ(1-4)GlcNAcβ(1-3)Galβ(1-4)GlcNAcβ(1-3)Galβ(1-4)GlcNAcβ(1-2)Manα(1-6)]-Manβ(1-4)GlcNAcβ(1-4)GlcNAcβ-(Lys-Val-Ala)Asn-Lys-Thr-NH2 | 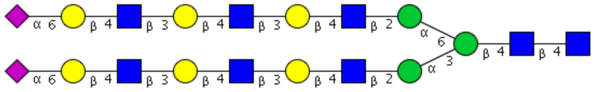 |
| **114** | NeuAcα(2-6)Galβ(1-4)GlcNAcβ(1-3)Galβ(1-4)GlcNAcβ(1-3)Galβ(1-4)GlcNAcβ(1-3)Galβ(1-4)GlcNAcβ(1-2)Manα(1-3)[NeuAcα(2-6)Galβ(1-4)GlcNAcβ(1-3)Galβ(1-4)GlcNAcβ(1-3)Galβ(1-4)GlcNAcβ(1-3)Galβ(1-4)GlcNAcβ(1-2)Manα(1-6)]-Manβ(1-4)GlcNAcβ(1-4)GlcNAcβ-(Lys-Val-Ala)Asn-Lys-Thr-NH2 | 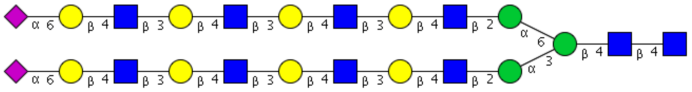 |
| **115** | NeuAcα(2-6)Galβ(1-4)GlcNAcβ(1-3)Galβ(1-4)GlcNAcβ(1-2)Manα(1-3)[NeuAcα(2-6)Galβ(1-4)GlcNAcβ(1-3)Galβ(1-4)GlcNAcβ(1-2)Manα(1-6)]-Manβ(1-4)GlcNAcβ(1-4)[Fucα(1-6)]-GlcNAcβ-(Lys-Val-Ala)Asn-Lys-Thr-NH2 | 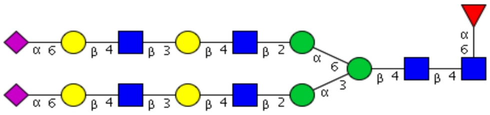 |
| **116** | NeuAcα(2-6)Galβ(1-4)GlcNAcβ(1-3)Galβ(1-4)GlcNAcβ(1-3)Galβ(1-4)GlcNAcβ(1-2)Manα(1-3)[NeuAcα(2-6)Galβ(1-4)GlcNAcβ(1-3)Galβ(1-4)GlcNAcβ(1-3)Galβ(1-4)GlcNAcβ(1-2)Manα(1-6)]-Manβ(1-4)GlcNAcβ(1-4)[Fucα(1-6)]-GlcNAcβ-(Lys-Val-Ala)Asn-Lys-Thr-NH2 | 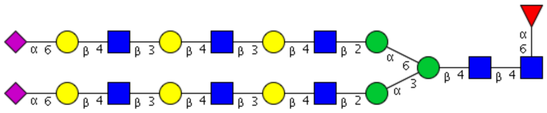 |

| Glycan # | Common Name | Structure |
|---|---|---|
| **117** | NeuAcα(2-6)Galβ(1-4)GlcNAcβ(1-3)Galβ(1-4)GlcNAcβ(1-3)Galβ(1-4)GlcNAcβ(1-3)Galβ(1-4)GlcNAcβ(1-2)Manα(1-3)[NeuAcα(2-6)Galβ(1-4)GlcNAcβ(1-3)Galβ(1-4)GlcNAcβ(1-3)Galβ(1-4)GlcNAcβ(1-3)Galβ(1-4)GlcNAcβ(1-2)Manα(1-6)]-Manβ(1-4)GlcNAcβ(1-4)[Fucα(1-6)]-GlcNAcβ-(Lys-Val-Ala)Asn-Lys-Thr-NH2 | 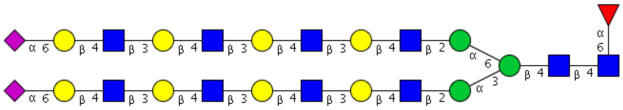 |
| **118** | NeuAcα(2-6)Galβ(1-4)GlcNAcβ(1-3)Galβ(1-4)GlcNAcβ(1-2)Manα(1-3){NeuAcα(2-6)Galβ(1-4)GlcNAcβ(1-3)Galβ(1-4)GlcNAcβ(1-2)[NeuAcα(2-6)Galβ(1-4)GlcNAcβ(1-3)Galβ(1-4)GlcNAcβ(1-6)Manα(1-6)]}-Manβ(1-4)GlcNAcβ(1-4)GlcNAcβ-(Lys-Val-Ala)Asn-Lys-Thr-NH2 | 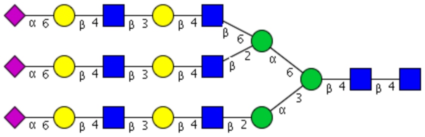 |
| **119** | NeuAcα(2-6)Galβ(1-4)GlcNAcβ(1-3)Galβ(1-4)GlcNAcβ(1-2)Manα(1-3){NeuAcα(2-6)Galβ(1-4)GlcNAcβ(1-3)Galβ(1-4)GlcNAcβ(1-2)[NeuAcα(2-6)Galβ(1-4)GlcNAcβ(1-3)Galβ(1-4)GlcNAcβ(1-6)Manα(1-6)]}-Manβ(1-4)GlcNAcβ(1-4)[Fucα(1-6)]-GlcNAcβ-(Lys-Val-Ala)Asn-Lys-Thr-NH2 | 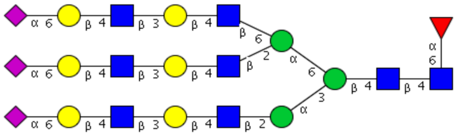 |
| **120** | LN/6'SLN/6'SLN-TriN | 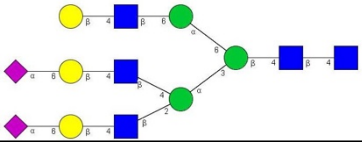 |
| **121** | 6'SLN/LeX/LeX-TriN | 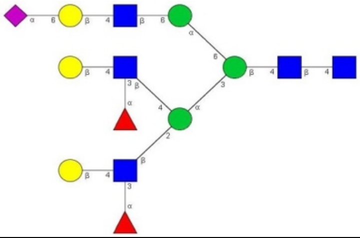 |
| **122** | 6'SLNLN/LeX/LeX-TriN | 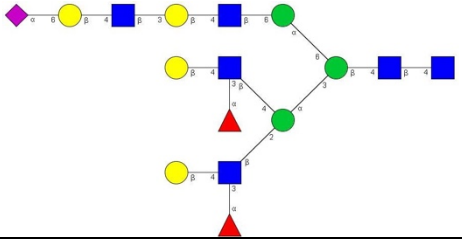 |
