## Supplementary material for "Sialic acid ligands of CD28 block co-stimulation of T cells": SI Data

### Supplementary Glycan Microarray Document

### Based on MIRAGE Guidelines (doi:10.3762/mirage.3)

| Classification | Guidelines |
|---|---|
| **1. Sample: Glycan Binding Sample** | |
| Description of Sample | Recombinant immune synapse proteins originating from human and mouse.<br><br>Figure 2C:<br><br>Human CD80<br>Human CD28<br>Human CD28 + CD80 (complex)<br>Mouse CD80<br>Mouse CD28<br>Mouse CD28 + CD80 (complex)<br><br>Extended Data Figure 8:<br><br>Human CD80 (duplicate from 2C)<br>Human CD86<br>Human CTLA4<br>Human PD-1<br>Human PD-L1<br>Human PD-L2<br>Mouse CD80 (duplicate from 2C)<br>Mouse CD86<br>Mouse CTLA4<br>Mouse PD-1<br>Mouse PD-L1<br>Mouse PD-L2<br><br>Full-length human and mouse CD80 genes were isolated, respectively, from cDNA libraries from human Daudi cells and ex vivo mouse splenocytes. Briefly, RNA extraction was performed on approximately 1 million cells (each) using the RNeasy Plus mini kit (Qiagen), according to the manufacturer's instructions. Purified total RNA was converted to a cDNA library via RT-PCR using ProtoScript II reverse transcriptase (NEB) and an 18-mer oligo-dT primer. Respective full-length CD80 genes were amplified using Q5 DNA polymerase (NEB) and the following primer sets: Human-FWD: 5'-ATG GGC CAC ACA CGG AGG CAG GGA ACA TCA C-3'; Human REV: 5'-TAC AGG GCG TAC ACT TTC CCT TCT CAA TCT CTC ATT CC-3'; Mouse FWD: ATG GCT TGC AAT TGT CAG TTG ATG CAG GAT ACA CC-3'; Mouse REV: 5'-AAG GAA GAC GGT CTG TTC AGC TAA TGC TTC TTC AGG-3'. Expression clones corresponding to the respective CD80 extracellular ectodomains (including Vset and Cset/C2) were amplified by PCR and subcloned via HiFi assembly (NEB) into a |

| | |
|---|---|
| | custom vector for mammalian cell expression encoding C-terminal eGFP and a $His_8$ tag using the following primer sets: Human-FWD: 5'-GGC TTC CGT CCT GGC AGG ATC AGT TAT CCA CGT GAC CAA G-3'; Human-REV: 5'-TTT CGC GCT TGA TCA GTG ATC CTG TAT TCC AGT TGA AGG T-3'; Mouse-FWD: 5'-GGC TTC CGT CCT GGC AGG ATC AGA ACA ACT GTC CAA GTC A-3'; Mouse-REV: 5'-TTT CGC GCT TGA TCA GTG ATC CTT TTT CCC AGG TGA AGT C-3'.<br><br>For expression, purified human and mouse CD80 plasmid stocks were transfected into 293F and CHO-K1 cell cultures, respectively, using pre-complexation with PEI-MAX (Polysciences) at a 4:1 mass:mass ratio (final). 293F cells expressing human CD80 were incubated at 37°C, 8% $CO_2$, 70% humidity for 5 days before harvesting condition media for purification. Adherent CHO-K1 cells were incubated in DNA:PEI mixture for 12 hours before exchanging into serum-free media and incubation for 4 days at 37°C, 5% $CO_2$. CD80-containing media were centrifuged at low speed to remove cells and large debris, vacuum-filtered over 0.8 µm nitrocellulose membranes, and applied directly to 5ml NiNTA FF crude columns (GE), preequilibrated in 30 mM HEPES pH 7.5, 300 mM NaCl, for $Ni^{2+}$-affinity purification. Bound CD80 was eluted in a 30 ml gradient, exchanging into 30 mM HEPES pH 7.5, 300 mM NaCl, 500 mM imidazole. Protein-containing fractions were identified and purity determined via SDS-PAGE analysis. Pure fractions were pooled, concentrated to approximately 1.0 mg $ml^{-1}$ (final), aliquoted, snap frozen in liquid $N_2$, and stored at -80°C.<br><br>For other samples, purified proteins were obtained commercially as recombinant human Fc chimeras (BioLegend). |
| Sample modifications | N/A |
| Assay protocol | Please see method section in the main text. |
| **2. Glycan Library** | |
| Glycan description for defined glycans | In-house sialoside array, consisting of 122 defined glycans (Extended Data Table 1). The synthesis of the contained glycans are described in Supplemental Experimental Procedures in (Peng et al., 2017). |
| Glycan description for undefined glycans | No glycans are undefined. |
| Glycan modifications | No modifications after initial synthesis were made. |
| **3. Printing Surface; e.g., Microarray Slide** | |
| Description of surface | NHS-ester functionalized hydro-polymer. |
| Manufacturer | Schott SlideH (Applied Microarrays 1070936). |
| Custom preparation of surface | None |

| | |
|---|---|
| Non-covalent Immobilization | All glycans are terminated with primary amine linker (either natural amino acid or chemical linker). |
| **4. Arrayer (Printer)** | |
| Description of Arrayer | MicroGrid II (Digilab) |
| Dispensing mechanism | Contact microarray pins (SMP3, ArrayIt) |
| Glycan deposition | Manufacturer estimation is 0.7nL per spot. However, actual delivery volume of each printed spot is not determined.<br>Each glycan was "pre-spotted" 3 times on Poly-L-Lysine derivatized slides (made in-house) before being spotted on SlideH slides. Each array contains 6 replicate spots of each individual glycan. |
| Printing conditions | Glycans were diluted to 100uM in 150mM NaPO4 buffer, pH 8.4 + 0.005% Tween-20. 10uL of each glycan was transferred to a 384-well microtiter plate and printed at ambient temperature and relative humidity of 50-65%. |
| **5. Glycan Microarray with "Map"** | |
| Array layout | Each slide contains 5 replicate arrays, consisting of a 4x2 (8) subarray pattern with each subarray containing 12x10 features (not all features contain a printed sample).<br>Array Layout file = "Sav7.GAL" |
| Glycan identification and quality control | In-house sialoside array, consisting of 122 defined glycans (Supplementary Table 1).<br>Quality control was assessed by incubation with plant lectins, AAL, ECA and SNA, to monitor fucosylations, de-sialylation and NeuAc-α2-6 terminated glycans, respectively. See Supplemental Experimental Procedures 2 in (Peng et al., 2017). |
| **6. Detector and Data Processing** | |
| Scanning hardware | Innoscan 1100AL (Innopsys) |
| Scanner settings | Scanning resolution: 10 µm / pixel<br>Laser channel: 488<br>PMT Voltages: Adjusted for each sample to achieve maximum signal without saturation of any single spot.<br>Scan power: Adjusted for each sample to achieve maximum signal without saturation of any single spot. |
| Image analysis software | Mapix (Innopsys) |
| Data processing | Output .txt files containing calculated data were processed in MS Excel to determine the mean signal value of 6 replicate spots with highest and lowest signals removed (e.g. average of 4 spots). |

| 7. Glycan Microarray Data Presentation | |
|---|---|
| Data presentation | The microarray binding results are in **Figure 2C,** and **Extended Data Figure 8**. Binding results are presented as 2D bar graphs with bars representing averaged mean signal of each glycan and error bars representing standard deviation. |
| **8. Interpretation and Conclusion from Microarray Data** | |
| Data interpretation | No software or algorithms were used to interpret processed data. |
| Conclusions | CD28 possesses a low-avidity secondary binding function as a sialoside-binding lectin. |
